# Biochemical regulation of Arabidopsis PUB33: a receptor-like cytoplasmic kinase with an integrated U-box domain that ubiquitinates *Ralstonia pseudosolanacearum* effector protein RipV1

**DOI:** 10.64898/2026.02.10.704836

**Authors:** Thakshila Dharmasena, Jihyun Choi, Injae Kim, Virginia N. Miguel, Natasha S. Kelkar, Maria Camila Rodriguez Gallo, Nicholas Hassan, Marco Trujillo, R. Glen Uhrig, Cécile Segonzac, Jacqueline Monaghan

## Abstract

Plant immunity relies on the detection of microbes and the rapid activation of intracellular defense pathways. Catalyzed by protein kinases and E3 ubiquitin ligases, respectively, phosphorylation and ubiquitination are among the most abundant post-translational modifications that regulate immune pathways. It has been well established that members of the receptor-like cytoplasmic kinase (RLCK) and plant U-box E3 ligase (PUB) families are critical components of plant immune signaling. Interestingly, a group of proteins that contain both an RLCK domain and a PUB domain has been conserved throughout plant evolution, referred to as subgroups RLCK-IXb and PUB-VI within their respective families. While very little is known about these proteins, evidence from multiple independent studies indicates that orthologous PUB-VI/RLCK-IXb proteins in potato, tomato, *Nicotiana benthamiana,* and *Arabidopsis thaliana* associate with diverse pathogen effectors from the oomycete pathogen *Phytophthora infestans*, bacterial pathogen *Ralstonia pseudosolanacearum,* and the mirid bug *Apolygus lucorum*, suggesting that they may be critical virulence targets or components of the immune response. However, the biochemical activities of these proteins and how they contribute to plant health remain poorly defined. Here, we introduce the PUB-VI/RLCK-IXb clade in Arabidopsis, focusing on PUB32, PUB33, and PUB50. We show that PUB33 exhibits dual kinase and E3 ubiquitin ligase activities that are inversely regulated by autophosphorylation at Thr333. PUB33 forms homomers and heteromers with PUB32 which attenuate PUB33 catalytic activity. Although we did not observe clear defects in innate immune signaling in *pub32, pub33*, or *pub50* mutants, we found that overexpression of *PUB33* can suppress cell death triggered by the *R. pseudosolanacearum* effector RipV1 in *N. benthamiana*. Moreover, PUB33 directly ubiquitinates RipV1 *in vitro* and reduces RipV1 accumulation *in planta,* suggesting that it functions as part of the immune response against *R. pseudosolanacearum*.

## Introduction

Post-translational modifications increase the diversity, complexity, and heterogeneity of proteomes by regulating protein conformation, function, localization, and interaction partners (Prabakaran et al. 2012). Among these modifications, phosphorylation and ubiquitination are the most abundant in eukaryotes (Nguyen et al. 2013). Phosphorylation is catalyzed by protein kinases that transfer the gamma phosphate moiety from ATP to specific serine, threonine, or tyrosine residues on protein substrates (Hunter 2007; Dievart et al. 2020). Comparatively, ubiquitination involves three enzymes (E1 ubiquitin activating enzyme, E2 ubiquitin conjugating enzyme, and E3 ubiquitin ligase enzyme) that work sequentially to covalently attach the 76 amino acid protein ubiquitin to target proteins, typically on Lys residues (Nguyen et al. 2013). Ubiquitin itself possesses seven Lys residues including K48 and K63 that can participate in poly-ubiquitin chain formation (Trenner et al. 2022). Substrates can be mono-ubiquitinated (i.e., the attachment of a single ubiquitin to one residue), multi-mono-ubiquitinated (i.e., the attachment of a single ubiquitin to multiple residues), or poly-ubiquitinated (i.e., the end-to-end attachment of linked ubiquitin proteins to at least one residue) (Nguyen et al. 2013). The type of ubiquitination directs the subsequent fate of the target protein (Trenner et al. 2022). For example, cytoplasmic and nuclear proteins modified with K48-linked polyubiquitin are typically marked for degradation via the 26S proteasome, while plasma membrane proteins modified with K63-linked polyubiquitin enter the endovacuolar degradation pathway (Trenner et al. 2022; Isono et al. 2024). Together, phosphorylation and ubiquitination play critical roles in regulating broad cellular processes such as the cell cycle, metabolic pathways, growth, transcriptional control, and stress signaling (Lehti-Shiu and Shiu 2012; Morreale and Walden 2016).

Plant U-box proteins (PUBs) are a large group of E3 ligases with documented roles in various stress and developmental pathways (Trenner et al. 2022). There are 64 PUBs in *Arabidopsis thaliana*, phylogenetically divided into 10 classes based on the presence of auxiliary domains (Trenner et al. 2022). The most-studied PUBs belong to class IV which contain an Armadillo repeat domain C-terminal to the U-box domain (Trenner et al. 2022). Intriguingly, class VI PUBs contain a receptor-like cytoplasmic kinase (RLCK) domain N-terminal to the U-box domain (Azevedo et al. 2001; Trenner et al. 2022). Class VI PUBs are thus also classified within the RLCK-IXb subfamily. Related to the Interleukin-1 associated kinase (IRAK) group of kinases, RLCKs belong to the plant receptor-like kinase (RLK) family comprised of over 600 genes divided into 64 subfamilies in *Arabidopsis* (Shiu and Bleecker 2001; Lehti-Shiu and Shiu 2012). RLCKs serve as key players in signal transduction of many stress signaling pathways (Liang and Zhou 2018). Prominent examples include BOTRYTIS INDUCED KINASE1 (BIK1) and related AvrPphB SUSCEPTIBLE 1-LIKE (PBLs), RLCKs from the VII subfamily that are critical to immune signaling (Lu et al. 2010; Zhang et al. 2010; Laluk et al. 2011; Rao et al. 2018; DeFalco and Zipfel 2021; Gonçalves Dias et al. 2022). Immune signaling is initiated at the plant cell surface, where pattern recognition receptors (PRRs) bind microbe-associated molecular patterns (MAMPs) such as flg22, the immunogenic peptide from flagellin (Chinchilla et al. 2007; Buscaill et al. 2024; Matsui et al. 2024). Most PRRs are integral membrane receptor kinases and MAMP binding to the ectodomain results in kinase activation (DeFalco and Zipfel 2021). BIK1 is a convergent substrate of multiple PRRs and represents a signaling bottleneck that is under tight regulatory control (DeFalco and Zipfel 2021; Gonçalves Dias et al. 2022). As part of their virulence strategy, pathogens secrete effector proteins into host cells to evade or suppress immune signaling (Toruño et al. 2016). Highlighting their importance to immune signaling, BIK1 and other RLCKs are the targets of multiple effectors from phylogenetically distant pathogens (Shao et al. 2003; Zhang et al. 2010; Feng et al. 2012; Guy et al. 2013; Blekemolen et al. 2023; Sun et al. 2023). In addition, BIK1 protein accumulation is directly regulated by class IV PUBs PUB22, PUB23, PUB24, PUB25, and PUB26 (Wang et al. 2018; Dou et al. 2025), as well as the class V PUB PUB4 (Yu et al. 2022). BIK1 is also monoubiquitinated by E3 ligases from another protein family that contributes to its activation status and subcellular localization (Ma et al. 2020).

Given the prominent roles of both PUBs and RLCKs in stress signaling, and the interplay between phosphorylation and ubiquitination on protein fates (Hunter 2007; Zhang and Zeng 2020; Gonçalves Dias et al. 2022), the presence of a group of proteins that contain both domains is intriguing. The PUB-VI/RLCK-IXb group does not contain KEEP ON GOING (KEG), a unique singleton protein that contains an E3 ligase domain and a protein kinase domain in addition to ankyrin repeats with broad roles in stress signaling (Stone et al. 2006; Zhou and Zeng 2017). Rather, the PUB-VI/RLCK-IXb group is estimated to have emerged approximately 1,500 million years ago, around the same time as canonical RLKs (Dievart et al. 2020). The conservation of the PUB-VI/RLCK-IXb group across species spanning evolutionary time implies critical functional roles. And yet, very little is known about these proteins. Of the nine PUB-VI/RLCK-IXb proteins encoded in Arabidopsis (**Figure 1A**), only PUB35 and PUB33 have been investigated. PUB35 negatively regulates abscisic acid (ABA) signaling and directly ubiquitinates the ABA transcription factor ABA INSENSITIVE 5 (ABI5) (Du et al. 2024). While the function of Arabidopsis PUB33 is largely unknown, putative orthologs of PUB33 in Solanaceous plants including potato (*Solanum tuberosum*), tomato (*Solanum lycopersicum*), and *Nicotiana benthamiana* are targeted by multiple effectors from diverse pathogens (He et al. 2019; McLellan et al. 2022; Dong et al. 2023; Choi et al. 2026), suggesting a role in plant immunity.

**Figure 1.**
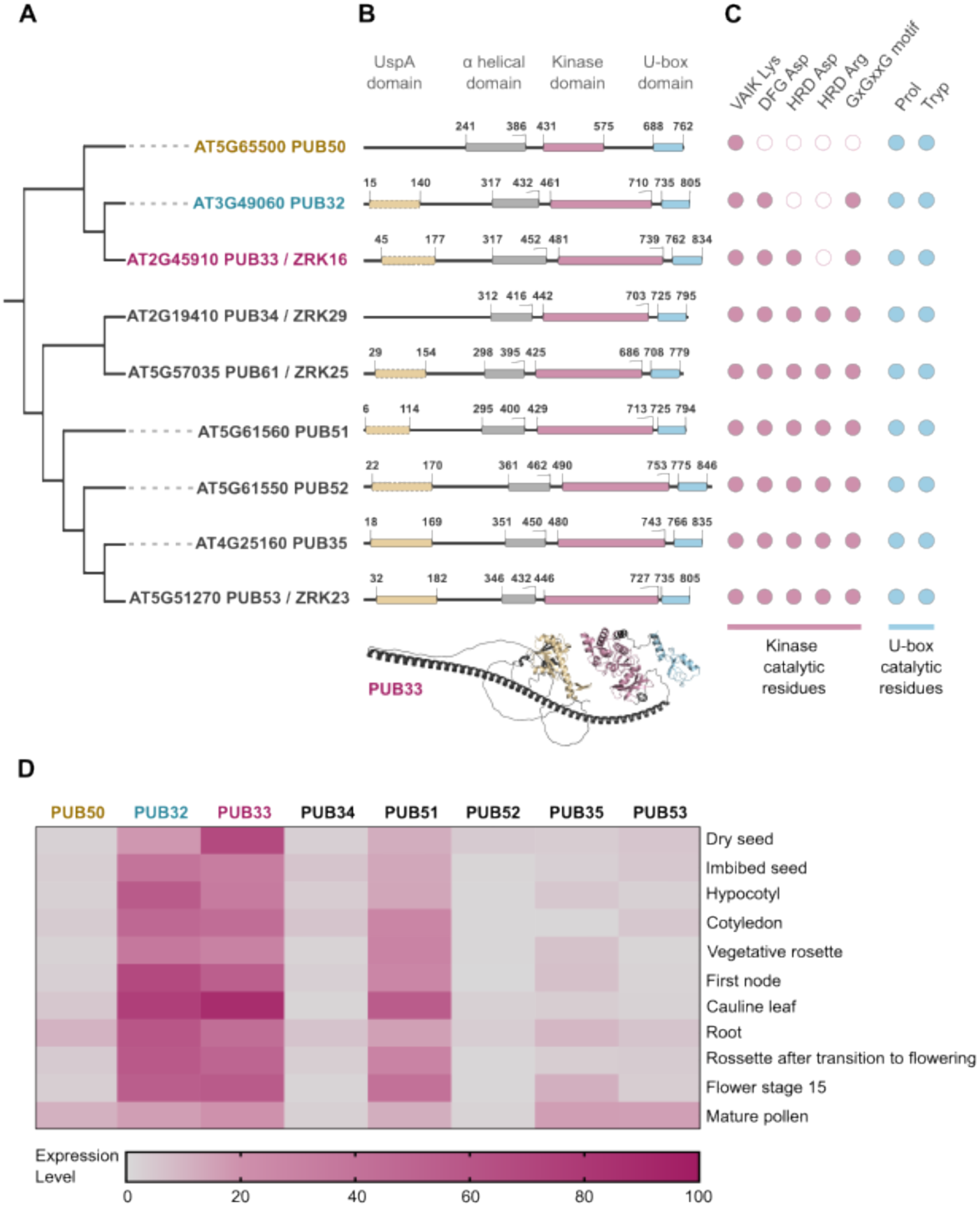
The Arabidopsis PUB-VI/RLCK-IXb subgroup contains 9 members. **(A)** Maximum-likelihood phylogenetic tree of *A. thaliana* RLCK-IXb proteins supported by 100 bootstraps. **(B)** Domain architecture of PUB-VI/RLCK-IXb proteins predicted by Pfam analysis (EMBL-EBI) and manual inspection of AlphaFold structural predictions. The UspA, α-helical, kinase, and U-box domains are shown in yellow, grey, pink, and blue, respectively. UspA domains with dashed lines indicate degeneration. The numbers delineate domain boundaries based on amino acid positions. The AlphaFold predicted structure for PUB33 is shown as an example for PUB-VI/RLCK-IXb proteins. Average predicted Local Distance Difference Test (pLDDT) score is 77.94. **(C)** *In silico* prediction of kinase and E3 ligase catalytic activity of RLCK-IXb proteins. The presence (closed pink circles) or absence (open pink circles) of conserved kinase residues include the ATP-binding Lys in the VAIK motif; the MgCl_2_ coordinating Asp in the DFG motif; the Asp residue critical for enzymatic activity in the HRD motif; the Arg residue that defines ‘RD’ or ‘non-RD’ kinases in the HRD motif; and the glycine-rich GxGxxG motif. The presence (closed blue circles) of conserved E2-binding Pro and Trp residues in the U-box domains are also shown. **(D)** Transcript expression profiles of PUB-VI/RLCK-IXb genes across developmental stages, as reported from global transcriptome studies (Nakabayashi et al. 2005; Schmid et al. 2005) and curated in the Arabidopsis eFP Brower (Winter et al. 2007). Values represent normalized microarray signal intensities (‘absolute’ expression levels) generated using GeneChip Operating Software with global scaling to a target signal value of 100. All analyses and visualization completed by TD.

The oomycete pathogen *Phytophthora infestans* secretes RXLR effectors (virulence factors containing a conserved RXLR motif, where X is any amino acid) into host cells during an infection (Wang et al. 2023). One of these effectors, PiSFI3/Pi06087/PexRD16, interacts with potato U-BOX-KINASE PROTEIN (StUBK) in the plant nucleus (He et al. 2019). Phylogenetics analysis indicates that StUBK is orthologous to AtPUB33 (He et al. 2019; Choi et al. 2026). While PiSFI13 suppresses immune responses and results in higher susceptibility to *P. infestans* when expressed in plant cells, over-expression of StUBK reduces this effect (He et al. 2019). Moreover, silencing StUBK or the *N. benthamiana* ortholog NbUBK results in higher colonization by *P. infestans*, while over-expression of NbUBK, StUBK, or AtPUB33 results in enhanced resistance to that pathogen in *N. benthamiana* or potato (He et al. 2019; McLellan et al. 2022). Arabidopsis is not a host of *P. infestans* and although PiSFI13 can bind NbUBK and StUBK, it is not able to bind AtPUB33, a feature which may be exploited to enhance resistance to *P. infestans* in potato (McLellan et al. 2022). In tomato, SlRLCK-IXb-1/SlPUB33 is ubiquitinated by RipV1, a novel E3 ligase (NEL) effector from the bacterial vascular pathogen *Ralstonia pseudosolanacearum* (Choi et al. 2026). Contrary to the PiSFI3-StUBK interaction which occurs in the nucleus, RipV1 likely interacts with SlRLCK-IXb-1 at the plant cell periphery, and may function to stabilize SlRLCK-IXb-1 (Choi et al. 2026). Co-expression with SlRLCK-IXb-1 suppresses RipV1-induced cell death in *N. benthamiana* (Choi et al. 2026), suggesting that SlRLCK-IXb-1 is either the target of RipV1 or is part of the plant defence arsenal against RipV1. Another RLCK-IXb protein in *N. benthamiana*, Niben261Scf01408g0000002/NbD000250, is targeted by the cyclophilin effector Al106 of the mirid bug *Apolygus lucorum* (Dong et al. 2023). This protein was referred to in that study as NbPUB33, however, phylogenetics analysis indicates that NbD000250 does not have a single putative ortholog in Arabidopsis and is rather found in a separate group that is basal to AtPUB32, AtPUB33, and AtPUB50 (Choi et al. 2026). As it is more closely related to SlRLCK-IXb-13, we will refer to it here as NbRLCK-IXb-13 to avoid confusion with NbUBK (He et al. 2019; McLellan et al. 2022). Al106 enhances plant susceptibility to biotic stresses including insect feeding and pathogen infection and interacts with both NbRLCK-IXb-13 and AtPUB33 in the cytoplasm (Dong et al. 2023). Al106 possesses peptidyl-prolyl-cis-trans-isomerase (PPIase) activity that is necessary for its ability to suppress the E3 ligase function of NbRLCK-IXb-13 *in vitro*, suggesting that it inhibits NbRLCK-IXb-13 function as part of its virulence strategy (Dong et al. 2023).

Although PUB33 orthologs in multiple species are targeted by effectors from diverse pathogens, the mechanism by which PUB33 contributes to plant immunity is not yet clear. While silencing *StUBK* in potato results in slightly lower expression of flg22-induced marker genes *StWRKY27* and *StACRE31* (He et al. 2019), silencing *NbUBK/NbRLCK-IXb-1* and *NbPUB32/NbRLCK-IXb-2* in *N. benthamiana* results in a higher flg22-induced burst of reactive oxygen species (ROS) (Choi et al. 2026). Furthermore, silencing *NbRLCK-IXb-13* in *N. benthamiana* results in enhanced susceptibility to cotton bullworm and *Phytophthora capsici* infections and a *pub33* mutant was reported to have reduced flg22-triggered ROS in Arabidopsis (Dong et al. 2023). In addition, RNAi-mediated silencing of GmPUB33A in soybean roots results in enhanced resistance to soybean cyst nematode (SCN) (Qi et al. 2022) and transcriptional reprogramming (Qi et al. 2025). Together, these results suggest that PUB33 may function in MAMP-triggered immune signaling with differing roles depending on the plant species.

Here, we provide an overview of Arabidopsis PUB-VI/RLCK-IXb proteins and characterize the biochemical properties of PUB32, PUB33, and PUB50, which form a distinct monophyletic group and are amongst the most highly expressed across developmental stages and tissue types. We find that only PUB33 exhibits both kinase and E3 ligase activities that are inversely regulated by autophosphorylation on residue Thr333. We present evidence that PUB33 is capable of forming homomers with itself and heteromers with PUB32, and that heteromer formation results in attenuated PUB33 catalytic activity. Although we did not observe defects in immune signaling in *pub33* mutants, we did observe that over-expression of *PUB33* suppresses cell death triggered by the *R. pseudosolanacearum* effector protein RipV1 in *N. benthamiana*. Furthermore, we find that PUB33 can ubiquitinate RipV1 *in vitro* and reduce RipV1 accumulation *in planta*. Our study provides key insights into the understudied but intriguing PUB-VI/RLCK-IXb proteins.

## Results & Discussion

### The PUB-VI/RLCK-IXb group contains 9 members in Arabidopsis

Proteins in the PUB-VI/RLCK-IXb group also contain a putative universal stress protein A (USP) domain and a long α-helical domain N-terminal to the RLCK and PUB domains (**Figure 1A**). In Arabidopsis, the proteins in this group include PUB32 (AT3G49060), PUB33/ZRK16 (AT2G45910), PUB34/ZRK29 (AT2G19410), PUB35 (AT4G25160), PUB50 (AT5G65500), PUB51 (AT5G61560), PUB52 (AT5G61550), PUB53/ZRK23 (AT5G51270), and PUB61/ZRK25 (AT5G57035) (Lewis et al. 2013; Trujillo 2018; Trenner et al. 2022). Note that the gene AT3G07370 was previously curated as PUB61 in some Arabidopsis databases but is now known only as CARBOXYL TERMINUS OF HSC70-INTERACTING PROTEIN (CHIP) and is not part of the PUB-VI/RLCK-IXb group. As members of the USP, PUB-VI, and RLCK-IXb families, we thought it prudent to determine their phylogenetic relationships within each of these families (**Supplementary Figure S1A-C).** As the nine members cluster similarly within USP, RLCK, and PUB families, we opted to use their RLCK phylogenetic classification as our reference (**Figure 1A**). While USPs are small proteins found across kingdoms with roles in stress tolerance, the function of USP domains within larger proteins is not well understood (Kerk et al. 2003; Chi et al. 2019). Protein family (Pfam) domain analysis using the European Molecular Biology Laboratory’s (EMBL) European Bioinformatics Institute (EBI) database suggests that many of the PUB-VI/RLCK-IXb proteins in Arabidopsis contain degenerate or low-confidence USP domains (**Figure 1B**). The RLCK-IXb subfamily contains 20 proteins (9 of which contain the U-box domain) and is phylogenetically related to the RLCK-XII-2 subfamily (**Supplemental Figure S1A**), which contains HopZ1 EFFECTOR TRIGGERED IMMUNITY DEFICIENT 1 (ZED1) and ZED1-related kinases (ZRKs) – pseudokinases that act as decoys or scaffolds to sense pathogen effectors (Lewis et al. 2013; Roux et al. 2014). Upon effector-mediated modification, these pseudokinases promote the formation of a signaling complex with RLCKs and nucleotide binding leucine-rich repeat (NLR) immune receptors such as HopZ-ACTIVATED RESISTANCE 1 (ZAR1), leading to assembly of a resistosome and activation of immune responses (Lewis et al. 2013; Wang et al. 2015, 2019; Seto et al. 2017; Gong et al. 2022). While most RLCK-IXb proteins containing a U-box domain are predicted to be active kinases, two are predicted to be pseudokinases (**Figure 1C**) as they lack the Asp in the HRD motif that is critical for catalytic activity (Taylor et al. 2021). While phylogenetically related, RLCK-XII-2 proteins do not contain the other domains present in PUB-VI/RLCK-IXb proteins and are thus likely to play distinct biological roles. Although no other PUB subfamily contains proteins that also harbor a kinase domain, class VII PUBs contain both the USP and α-helical domains N-terminal to the U-box and include PUB36 (AT3G61390), PUB37 (AT2G45920), PUB54 (AT1G01680), PUB55 (AT1G01660), and PUB56 (AT1G01670) (Trenner et al. 2022) (**Supplemental Figure S1B,D**). As class VII PUBs are only found in Brassicaceae, it has been proposed that they lost the kinase domain through evolution (Trenner et al. 2022). It is worth noting that all PUB-VI/RLCK-IXb proteins contain canonical U-box domains and are predicted to be active E3 ligases (**Figure 1C**). To gain some insight into their biological functions, we mined ePlant (Waese et al. 2017) for PUB-VI/RLCK-IXb expression patterns across developmental phases and tissue types (**Figure 1D**). The available data indicates that *PUB32*, *PUB33*, and *PUB51* are the most highly and broadly expressed, while the other loci are expressed during specific growth stages or tissues. As PUB32, PUB33, and PUB50 are putative homologs (**Figure 1A**) that are among the most highly expressed (**Figure 1D**) with potential roles in plant immune signaling (He et al. 2019; McLellan et al. 2022; Dong et al. 2023; Choi et al. 2026), we chose to study them further. Moreover, PUB32 and PUB50 are the only members of the PUB-VI/RLCK-IXb group unlikely to harbour kinase activity (**Figure 1C**), allowing for a comparative study to understand the dual functionality of kinase and E3 ligase activities in this intriguing group of proteins.

### PUB33 is both an active E3 ubiquitin ligase and an active protein kinase

The U-box domains in PUB32, PUB33, and PUB50 possess conserved Pro and Trp residues that are critical for binding E2 conjugating enzymes (**Figure 1C**) (Trujillo 2018) and are therefore expected to be catalytically active. Auto-ubiquitination is a common feature of PUBs (Trenner et al. 2022) and is visible as a smear of higher molecular weight proteins when all three enzymes and ubiquitin are present. To test their activity as E3 ligases, we purified full-length (FL) MBP-PUB32^FL^, His_6_-MBP-PUB33^FL^, and MBP-PUB50^FL^ proteins as N-terminal translational fusions with maltose binding protein (MBP). Alongside, we also purified variants mutated in residues required for E2 binding (PUB32^FL-W768A^, PUB33^FL-W796A^) and truncated variants lacking the entire U-box domains (PUB32^ΔUBOX^, PUB33^ΔUBOX^, PUB50^ΔUBOX^) to serve as assay controls. When incubated *in vitro* with ubiquitin activating enzyme 1 (UBA1) and ubiquitin conjugating enzyme 8 (UBC8) we could readily detect auto-ubiquitination for PUB33 and PUB50, but surprisingly, not for PUB32 (**Figure 2A-C**). We conclude that only PUB33^FL^ and PUB50^FL^ are capable of auto-ubiquitination *in vitro*.

**Figure 2.**
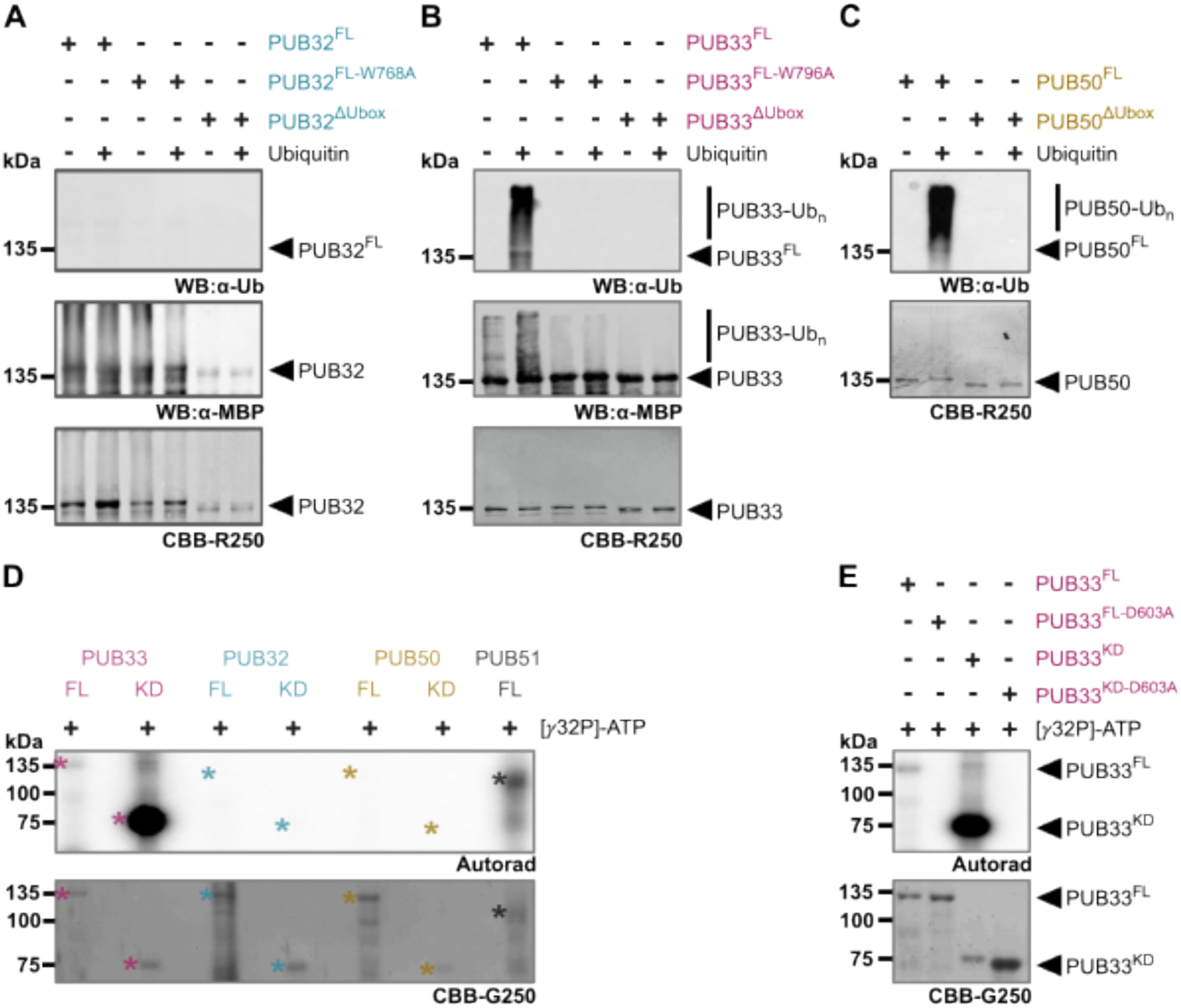
Analysis of PUB32, PUB33, and PUB50 *in vitro* catalytic activities. **(A–C)** *In vitro* auto-ubiquitination assays using **(A)** MBP-PUB32^FL^ compared to catalytic mutants MBP-PUB32^FL-W768A^ and MBP-PUB32^ΔUbox^; **(B)** His_6_-MBP-PUB33^FL^ compared to His_6_-MBP-PUB33^FL-W796A^ and His_6_-MBP-PUB33^ΔUbox^; **(C)** MBP-PUB50^FL^ compared to MBP-PUB50^ΔUbox^. Reaction mixtures included ubiquitin, ubiquitin activating protein 1 (UBA1) and ubiquitin conjugating protein 8 (UBC8). Western blots were probed with antibodies against ubiquitin (anti-Ub) or maltose binding protein (anti-MBP); protein loading is indicated by post-staining with Coomassie Brilliant Blue (CBB). **(D-E)** *In vitro* auto-phosphorylation assays of His_6_-MBP-PUB33^FL^, His_6_-MBP-PUB33^KD^, MBP-PUB32^FL^, MBP-PUB32^KD^, MBP-PUB50^FL^, MBP-PUB50^KD^, and His_6_-MBP-PUB51^FL^ (**D**) and His_6_-MBP-PUB33^FL^, MBP-PUB33^FL-D603A^, His_6_-MBP-PUB33^KD^, and MBP-PUB33^KD-D603A^ (**E**). Autoradiographs (Autorad) show incorporation of γ-³²P, and protein loading is indicated by post-staining with Coomassie Brilliant Blue (CBB). Asterisks indicate the respective proteins. All assays were repeated at least three times with similar results by TD.

Because PUB50 contains a truncated kinase domain and PUB32 is categorized as a pseudokinase (Kwon et al. 2019; Paul and Srinivasan 2020), only PUB33 is predicted to be an active kinase. Indeed, *in vitro* kinase assays using radioactive γ^32^P-ATP indicated that only PUB33^FL^ possesses auto-phosphorylation activity (**Figure 2D**). Auto-phosphorylation is a common feature of RLCKs (Liang and Zhou 2018; Hailemariam et al. 2024), visible by the incorporation of the radiolabelled γ phosphate of ATP onto residues of the kinase itself. Notably, PUB33 is the only PUB-VI/RLCK-IXb protein characterized as a non-RD kinase, as it lacks an Arg in the HRD motif located in the activation loop (**Figure 1C**). Non-RD kinases usually display reduced auto-phosphorylation and are a prevalent feature of receptor kinases involved in immune signaling (Dardick and Ronald 2006). We therefore also tested His_6_-MBP-PUB51^FL^ as an RD kinase (**Figure 1C**), and found that PUB51^FL^ demonstrated higher auto-catalytic activity than PUB33^FL^ (**Figure 2D**). Isolation of the kinase domains (KD) of PUB33, PUB32, and PUB50 displayed a similar trend as the FL proteins, however PUB33^KD^ possessed markedly higher auto-phosphorylation activity compared to PUB33^FL^ (**Figure 2D**). Importantly, a mutant variant of PUB33 with the catalytic Asp mutated to Ala, PUB33^KD-D603A^ lost its ability to auto-phosphorylate (**Figure 2E**). Overall, we conclude that PUB33 and PUB51 are active kinases, while PUB32 and PUB50 are not.

To our knowledge, the only other PUB-VI/RLCK-IXb protein that has been assessed for kinase function is SlRLCK-IXb-1. However, although predicted to be active, we could not detect any autophosphorylation using either SlRLCK-IXb-1^FL^ or SlRLCK-IXb-1^KD^ (Choi et al. 2026), leading to the hypothesis that the kinase domain of SlRLCK-IXb-1 may require cellular cues for activation. Alternatively, as an non-RD kinase, it may possess auto-catalytic activity that is below experimental detection *in vitro*. While PUB35 has been shown to ubiquitinate ABI5 (Du et al. 2024), its ability to phosphorylate ABI5 has not yet been assessed, nor has the importance of kinase capability to PUB35 function in ABA signaling. This is particularly relevant as the phospho-status of two SnRK-mediated phosphorylation sites on ABI5 impacts its ability to interact with PUB35 (Du et al. 2024). It will be important to test kinase and E3 ligase activities for other PUB-VI/RLCK-IXb proteins. For those that possess both catalytic activities, it will be important to decipher their *bone fide* phosphorylation and ubiquitination substrates. Do they catalyze both modifications onto target proteins? If so, are they sequential and do they influence each other? For example, a substrate may need to be phosphorylated prior to being ubiquitinated. Kinases and E3 ligases interact with their substrates in different ways, so if they do target the same substrate, it will be important to understand how their structural conformation coordinates that. Alternatively, or perhaps in tandem, the E3 ligase function of PUB-VI/RLCK-IXb proteins may be intrinsically regulated by auto-phosphorylation, or the kinase function may be regulated by auto-ubiquitination.

### Intra- or intermolecular inhibition of the PUB33 kinase domain

Our finding that PUB33^KD^ possessed higher auto-catalytic activity than PUB33^FL^ led to the hypothesis that the kinase domain may be subject to intramolecular regulation. The PUB33^KD^ variant contains the kinase domain as well as the 62 amino acid linker region between the kinase and U-box domains (**Figure 3A**). To assess if the N-terminal USP and α-helical domains inhibit the kinase domain, we compared PUB33^KD^, PUB33^FL^ and PUB33^ΔUBOX^ auto-phosphorylation *in vitro*, and observed similarly low levels of auto-phosphorylation on PUB33^FL^ and PUB33^ΔUBOX^ compared to PUB33^KD^ (**Figure 3B**). The PUB33^ΔUBOX^ variant also contains the linker region between the kinase and U-box, but the U-box domain is replaced by 65 other amino acids from the vector. These results suggest that the N-terminal USP and α-helical domains inhibit auto-phosphorylation by the kinase domain. We next generated PUB33^CT^, a variant excluding the entire N-terminal domain and containing only the C-terminal (CT) kinase, linker, and U-box (**Figure 3A**). We found that the PUB33^CT^ variant completely lost auto-phosphorylation activity (**Figure 3B**), suggesting that the U-box domain also suppresses the kinase domain. Furthermore, we also found that PUB33^FL-W796A^ displays reduced levels of auto-phosphorylation compared to PUB33^FL^ (**Figure 3C**), indicating that either auto-ubiquitination or the integrity of the U-box domain may be important for kinase activity. To test if engaging an E2 could release this inhibition, we performed *in vitro* auto-phosphorylation assays with PUB33^FL^ or PUB33^CT^ together with UBA1, UBC8, and ubiquitin. However, we could not detect auto-phosphorylation of either PUB33^FL^ or PUB33^CT^ when undergoing auto-ubiquitination (**Figure 3D**), suggesting that ubiquitination and phosphorylation cannot occur simultaneously.

**Figure 3.**
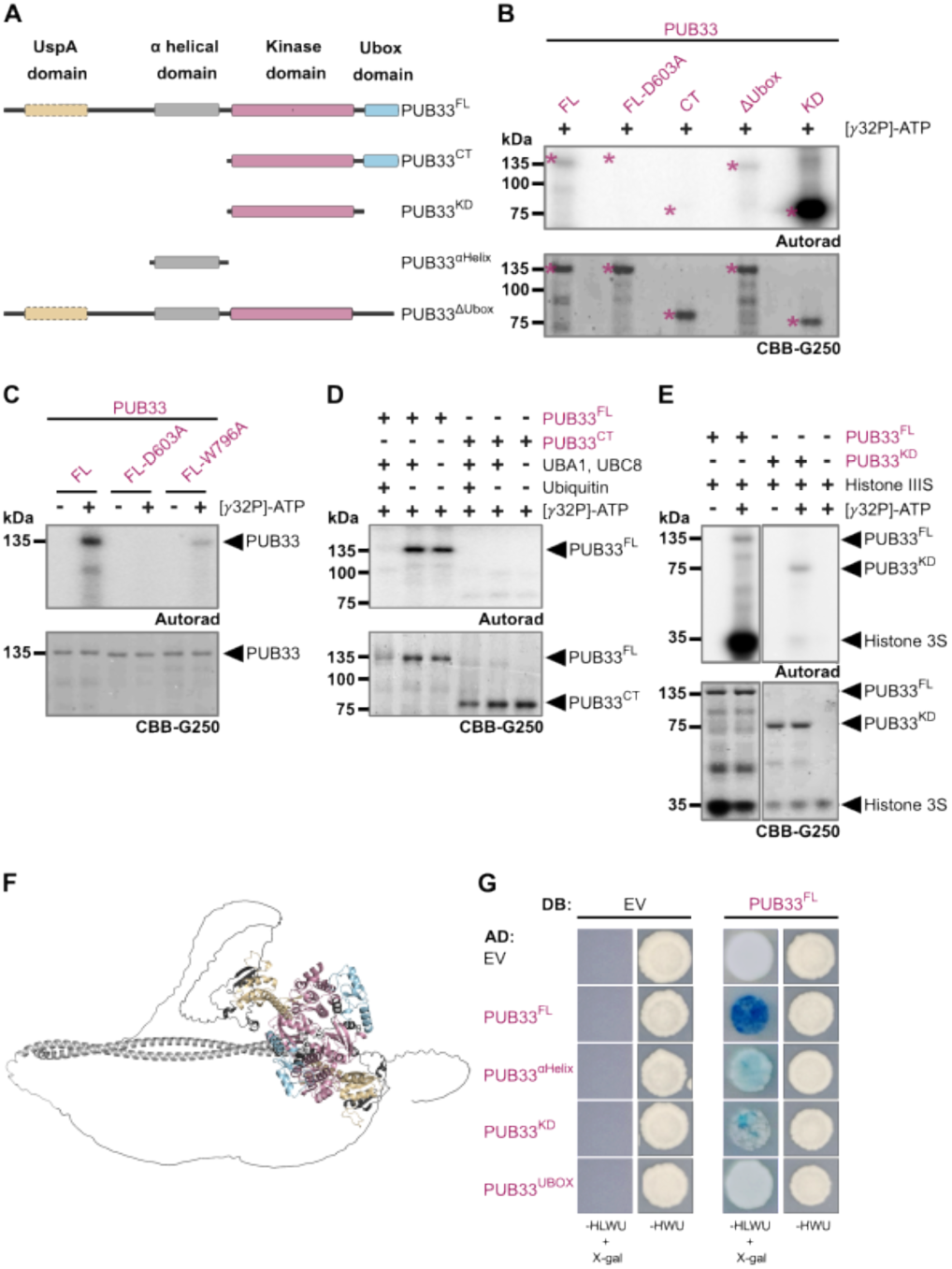
Inter- or intramolecular regulation of the PUB33 kinase domain. **(A)** Schematic representation of PUB33 protein domains and truncated variants: PUB33^FL^ indicates the full-length protein; PUB33^CT^ indicates the kinase and U-box domains and removal of the entire N-terminus; PUB33^ΔUbox^ indicates removal of the U-box domain; PUB33^KD^ indicates only the kinase domain; PUB33^αHelix^ indicates only the α-helical domain. Exact amino acid boundaries for all variants can be found in **Supplemental Table S2**. Schematic created by TD. **(B-D)** *In vitro* autophosphorylation assays of PUB33 variants as indicated. Autoradiography (Autorad) indicates γ-³²P incorporation, and protein loading is indicated by post-staining with Coomassie Brilliant Blue (CBB). Asterisks in B indicate expected proteins of interest. Note that the first and fourth lanes of the CBB-stained gel in **D** display characteristic ubiquitination smears corresponding to PUB33^FL^ or PUB33^CT,^, indicating that the ubiquitin ligase is active. Assays conducted at least three times by TD with similar results. **(E)** *In vitro* trans-phosphorylation assay between His_6_-MBP-PUB33^FL^ and His_6_-MBP-PUB33^KD^ and the universal substrate histone H3S. Autorad indicates γ-³²P incorporation, and protein loading is indicated by post-staining with CBB. Note that the signal from H3S is so strong that it masks the signal from His_6_-MBP-PUB33^KD^ when imaged together. Boxes indicate that some lanes were removed digitally for clarity but everything shown is from a single gel. Assays conducted at least three times by TD with similar results. **(F)** Possible configuration of a PUB33 homomer, as predicted by AlphaFold2. The colours match the labels in A. Average predicted Local Distance Difference Test (pLDDT) score for the PUB33 monomer is is 66.52 and the predicted template modeling (pTM) and interface pTM (ipTM) scores for the PUB33 homomer are 0.47 and 0.45, indicating low-to-moderate confidence in both the overall model and the predicted interface. Analysis by TD. **(G)** Yeast 2-hybrid between the indicated PUB33 variants. EV is the empty vector control. Growth was assessed on SD media lacking His, Leu, Trp, and Uracil (-HLWU) with the addition of X-galactosidase (X-gal) as markers for protein:protein interaction or on SD-HWU as a transformation control. DB indicates constructs translationally fused to the Locus for X-Ray Sensitivity A (LexA) DNA binding domain; AD indicates constructs translationally fused to the B42 transcriptional activation domain. Assays conducted by IK at least three times with similar results.

While auto-phosphorylation is widely documented to be required for full activation of many kinases, the biochemical mechanism must differ from that of trans-phosphorylation on a substrate protein (Beenstock et al. 2016). To test the trans-phosphorylation capability of PUB33, we assessed whether it is able to phosphorylate Histone III (H3) - a highly phosphorylatable protein that can be used as a general substrate when a biologically relevant substrate is unknown. Interestingly, although PUB33^FL^ has much lower auto-catalytic activity than PUB33^KD^ (**Figure 2E**), we found that it could readily transphosphorylate H3 (**Figure 3E**). Comparatively, and to our surprise, we found that PUB33^KD^ could not accept H3 as a substrate (**Figure 3E**). These data suggest that PUB33 has intrinsically high kinase activity that is regulated by intra- and/or inter-molecular mechanisms. We propose that the N-terminal region suppresses auto-phosphorylation while promoting a conformational state capable of substrate trans-phosphorylation. The U-box domain can also strongly repress kinase activity when isolated from the N-terminus, suggesting layered intramolecular control. Notably, deletion of the U-box in PUB33^ΔUBOX^ does not alter auto-catalysis in the context of full-length protein, indicating that these regulatory elements may function in distinct conformational contexts. We propose a model in which PUB33 toggles between auto-phosphorylation and substrate-phosphorylation states, potentially including additional induced activation conformations *in vivo*.

Dimerization is a common feature of U-box proteins (Trenner et al. 2022). AlphaFold2 predicts that the α-helical domains of two PUB33 proteins may homodimerize in a coiled-coil formation (**Figure 3F**). Although the confidence of this prediction is in the low-to-moderate range, and it is difficult to predict how the UspA, kinase, and U-box domains may interact in a cellular context, the suggestion of a coiled-coil interface between the α-helical domains was intriguing. To test if PUB33 can associate with itself, we generated a series of constructs representing the domains of PUB33 translationally fused to the Locus for X-Ray Sensitivity A (LexA) DNA-binding domain (DB) or the B42 transcriptional activation domain (AD) and conducted yeast 2-hybrid (Y2H). Because we observed some auto-activation of the leucine biosynthesis marker gene when DB-PUB33 variants were expressed alone (**Figure 3G**), we relied on the activation of β-galactosidase for our analysis. When expressed together, DB-PUB33^FL^ and PUB33^FL^-AD enabled yeast to produce blue precipitate in the presence of 5-bromo-4-chloro-3-indolyl-β-D-galactopyranoside (X-gal) (**Figure 3G**), indicating transcriptional activation of β-galactosidase. We also observed some minimal activation of β-galactosidase between DB-PUB33^FL^ and PUB33^α-helix^-AD or PUB33^KD^-AD (**Figure 3G**). These results indicate that PUB33 is capable of interacting with itself - either intramolecularly or intermolecularly as an oligomer. It is noteworthy that the FL proteins interacted much more strongly than any of the truncation constructs, indicating that the integrity of the entire protein is important for oligomerization or that there are multiple oligomerization surfaces that act actively or synergistically. It is also possible that the formation of DB-PUB33^FL^ homomers may outcompete any other interaction.

### PUB33 autophosphorylation on residue T333 modulates both its kinase and E3 ligase activity

PUBs are well known to be regulated by phosphorylation (Furlan et al. 2017; Wang et al. 2018; Zhou et al. 2018; Trenner et al. 2022). To assess if auto-phosphorylation regulates PUB33 E3 ligase function, we conducted *in vitro* auto-ubiquitination assays using UBA1, UBC8, and either the wild-type PUB33^FL^ or kinase-inactive PUB33^FL-D603A^ variant. We found that PUB33^FL-D603A^ exhibited strongly reduced auto-ubiquitination over a 30 min time course (**Figure 4A**). This result indicates that kinase activity regulates E3 ligase activity, either directly via auto-phosphorylation or indirectly via the conformation conferred by its active form. To map PUB33 auto-phosphorylation sites, we performed phosphoproteomics following an *in vitro* kinase assay using PUB33^FL^. We identified a total of four phosphorylation sites: three (T638, T641, and T643) clustered within the activation loop of the kinase, and one (T333) located at the N-terminus of the α-helical domain (**Figure 4B**). As the three phosphosites located in the activation loop are likely to directly contribute to the kinase function of PUB33, we aimed to further characterize the role of phosphorylation on T333, which is uniquely found in PUB33 (**Figure 4C**). Remarkably, we found that replacing Thr333 with Ala (PUB33^FL-T333A^) resulted in very strong reduction of E3 ligase function *in vitro* (**Figure 4D**), suggesting that phosphorylation of T333 is required for full E3 ligase activity. Alongside, we also replaced T333 with Asp (PUB33^FL-T333D^) to potentially mimic the negative charge conferred by phosphorylation at this site, however we found that PUB33^FL-T333D^ also resulted in a loss of E3 ligase function (**Figure 4D**), indicating that both T333A and T333D affect auto-ubiquitination. In addition, PUB33^CT^ displayed enhanced auto-ubiquitination compared to PUB33^FL^ (**Figure 4E**), providing further evidence that the E3 ligase domain is under negative regulatory control by the N-terminal domain.

**Figure 4.**
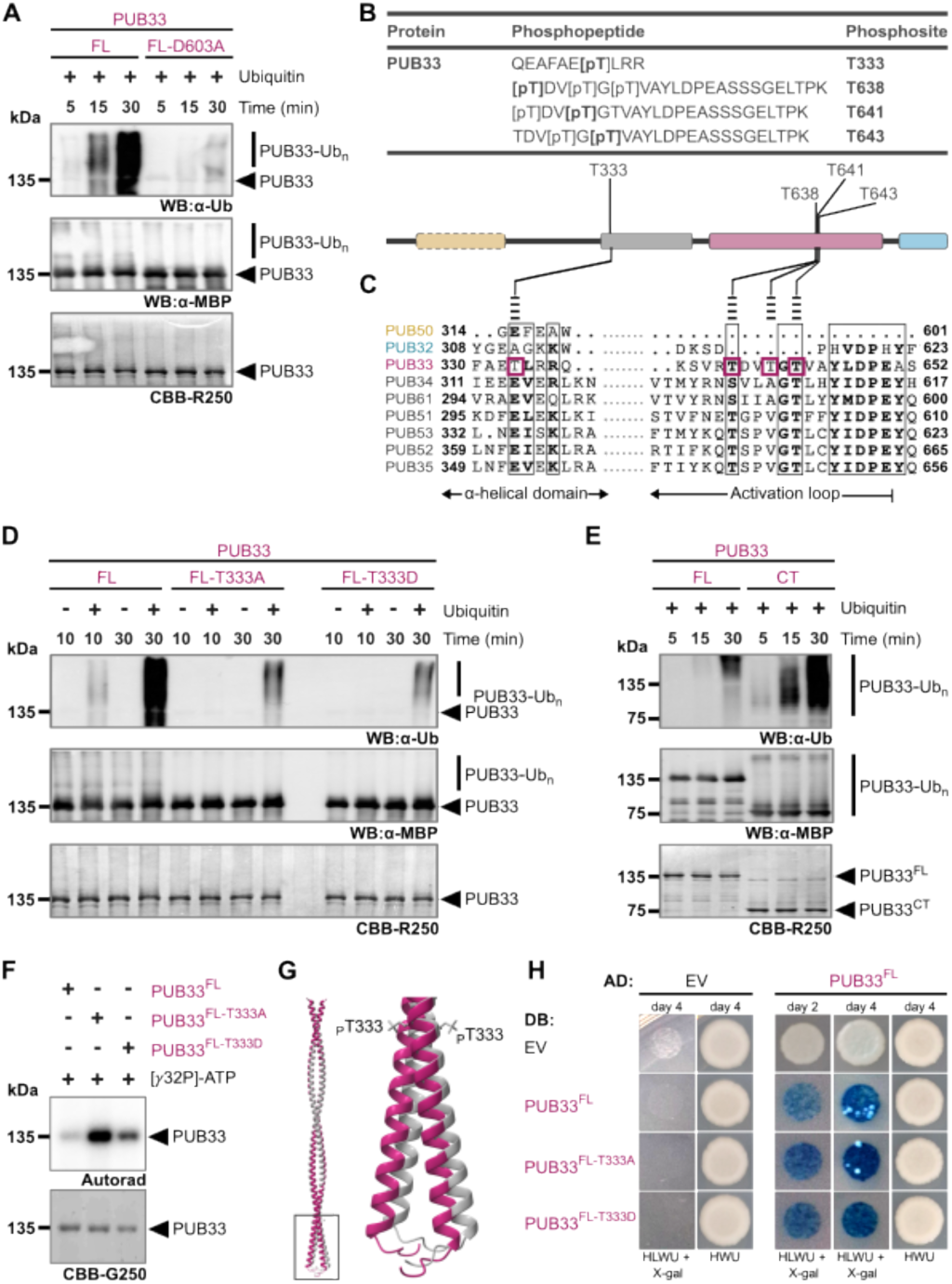
PUB33 autophosphorylation on residue T333 modulates both its kinase and E3 ligase activity. **(A)** *In vitro* auto-ubiquitination assays comparing His_6_-MBP-PUB33^FL^ with MBP-PUB33^FL-D603A^. The reaction mixture included ubiquitin, ubiquitin activating protein 1 (UBA1) or ubiquitin conjugating protein 8 (UBC8). Western blots were probed with antibodies against ubiquitin (anti-Ub) and maltose-binding protein (anti-MBP) and protein loading is indicated by post-staining with Coomassie Brilliant Blue (CBB). These assays were repeated more than three times with similar results by TD. **(B)** PUB33 phosphopeptides detected following *in vitro* His_6_-MBP-PUB33^FL^ autophosphorylation. Each peptide was identified in at least 2/3 independent replicates and was absent in control samples without ATP. Phosphosites are indicated in bold, with positions shown on the right. The schematic below indicates their location on PUB33^FL^; colours indicate the domains of PUB33 as defined in Figure 1. Kinase assays were performed by TD; trypsin digestion and LC-MS/MS analysis were performed by MCRG. **(C)** Multiple sequence alignment of the regions including the PUB33 phosphosites across the PUB-VI/RLCK-IXb proteins. Analysis by TD. **(D-E)** *In vitro* auto-ubiquitination assays comparing His_6_-MBP-PUB33^FL^ with the indicated variants. Reaction mixtures include UBA1 and UBC8, with (+) and without (-) ubiquitin for the indicated times. Western blots were probed with anti-Ub and anti-MBP, and protein loading is indicated by post-staining with CBB. These assays were repeated more than three times with similar results by TD. **(F)** *In vitro* auto-phosphorylation assays comparing His_6_-MBP-PUB33^FL^ to PUB33^FL-T333A^ and PUB33^FL-T333D^. Autoradiography (Autorad) indicates γ-³²P incorporation, and protein loading is indicated by post-staining with CBB. These assays were repeated more than three times with similar results by TD. **(G)** Alignment of the predicted coiled-coil domain of the PUB33 homomer in its unphosphorylated (magenta) and phosphorylated at T333 (grey) configurations, as predicted by AlphaFold3. Although we noticed a modest shift in this region, the predicted PUB33 and PUB33-pT333 homomers are excellently aligned overall, with a sequence alignment score of 2264.2 and a root mean square deviation of 0.713 Å. Analysis by JM. **(H)** Yeast 2-hybrid between the indicated PUB33 variants. EV is the empty vector control. Growth was assessed on SD media lacking His, Leu, Trp, and Uracil (-HLWU) with the addition of X-galactosidase (X-gal) as markers for protein:protein interaction or on SD-HWU as a transformation control. Assays conducted by IK at least three times with similar results.

Importantly, both PUB33^FL-T333A^ and PUB33^FL-T333D^ maintained kinase activity *in vitro* (**Figure 4F**), indicating that the impact of this mutation on E3 ligase function is not due to a lack of kinase function. In fact, we observed higher auto-catalytic activity in both PUB33^FL-T333A^ and PUB33^FL-T333D^ relative to PUB33^FL^ (**Figure 4F**), suggesting that phosphorylation at this site inhibits the kinase function of PUB33. Because we previously found that the N-terminal domain inhibits auto-catalysis of the PUB33 kinase domain, we reasoned that phosphorylation at T333 may impact PUB33 homomer formation. We used AlphaFold3 to model how phosphorylation at T333 may impact oligomerization, and noted a modest shift at the far N-terminus of the putative coiled-coil region (**Figure 4G**). Because the N-termini of the α-helical domains are proximal to very long regions of intrinsic disorder, it is difficult to interpret how this shift would impact oligomerization overall. We thus assessed if the PUB33^FL-T333A^ and PUB33^FL-T333D^ variants could interact with PUB33^FL^ in Y2H. We found that both PUB33^FL-T333A^ and PUB33^FL-T333D^ interacted more strongly with PUB33^FL^ (**Figure 4H**), suggesting that phosphorylation at this site may disrupt oligomerization. Given the possible geometry of the PUB33 homomer, we suggest that trans-autophosphorylation at T333 results in a conformational shift via the coiled-coil that promotes full E3 ligase functionality while simultaneously inhibiting kinase activity. This would mean that a ‘tighter’ homomer contributes to higher kinase activity and lower E3 ligase activity, while a ‘loosened’ homomer following phosphorylation of Thr333 results in lower kinase activity and higher E3 ligase activity.

### PUB32 inhibits the catalytic activity of PUB33

To gain some insight into the biological role of PUB33, we aimed to identify *in vivo* interaction partners. We therefore generated homozygous transgenic lines in Col-0 expressing PUB33 C-terminally tagged with green fluorescent protein (Col-0/35S:PUB33-GFP). We immunopurified PUB33-GFP from one of these lines, Col-0/35S:PUB33-GFP line #9-4, and performed tandem mass spectrometry. As a control, we used proteins similarly immunopurified from Col-0. We considered any peptides uniquely found in PUB33-GFP samples in at least 2 of 3 independent experimental replicates, as well as any peptides that were statistically more abundant in the PUB33-GFP samples according to SAINTq analysis (Teo et al. 2016), to be putative PUB33 association partners (**Supplementary Table S1)**. In total, we identified 104 putative binding partners of PUB33. While future work will be needed to decipher *bone fide* interactors and their functions, preliminary gene ontology (GO) term analysis indicated an enrichment for biological and molecular function terms associated with translation (‘translation’ *p* = 6.09^-4^; ‘structural constituent of ribosome’ *p* = 1.087^-4^; ‘mRNA binding’ *p* = 5.03^-8^) and metabolism (‘cellular oxidant detoxification *p* = 2.20^-4^; ‘peroxidase activity’ *p* = 7.96^-4^).

Aside from this, one protein that caught our attention was PUB32 (**Supplementary Table S1; Figure 5A**). We confirmed that PUB32 and PUB33 are capable of interacting with each other in Y2H (**Figure 5B**), however we did not observe interaction between two PUB32 proteins (**Figure 5B**). These results suggest that PUB32:PUB33 heteromers and PUB33 homomers can form, whereas PUB32 homomers may not. Interestingly, modeling predictions using AlphaFold2 suggest that the α-helical domains of PUB32 and PUB33 may be capable of interacting in a coiled-coil fashion (**Figure 5C**). This led to the idea that PUB32:PUB33 heteromers may function differently than PUB33 homomers. To get initial insight into this, we first tested if co-incubation with PUB32 could modulate the E3 ligase or kinase activities of PUB33. We conducted *in vitro* ubiquitination and kinase assays with PUB33^FL^ in the presence of increasing amounts of PUB32^FL^. We found that PUB32^FL^ reduced auto-ubiquitination (**Figure 5D**) and auto-phosphorylation of PUB33^FL^ (**Figure 5E**) in a dose-dependent manner, suggesting that PUB32:PUB33 heteromers repress PUB33 activity. Furthermore, we found that PUB32^FL^ interacted less with PUB33^FL-T333A^ and PUB33^FL-T333D^ in Y2H **(Figure 5F**), suggesting that phosphorylation at Thr333 may loosen PUB33 homomers and allow for heteromerization with PUB32 as a form of regulatory control. All together, these data suggest that PUB32 directly regulates PUB33 by modulating both its kinase and E3 ligase activities.

**Figure 5.**
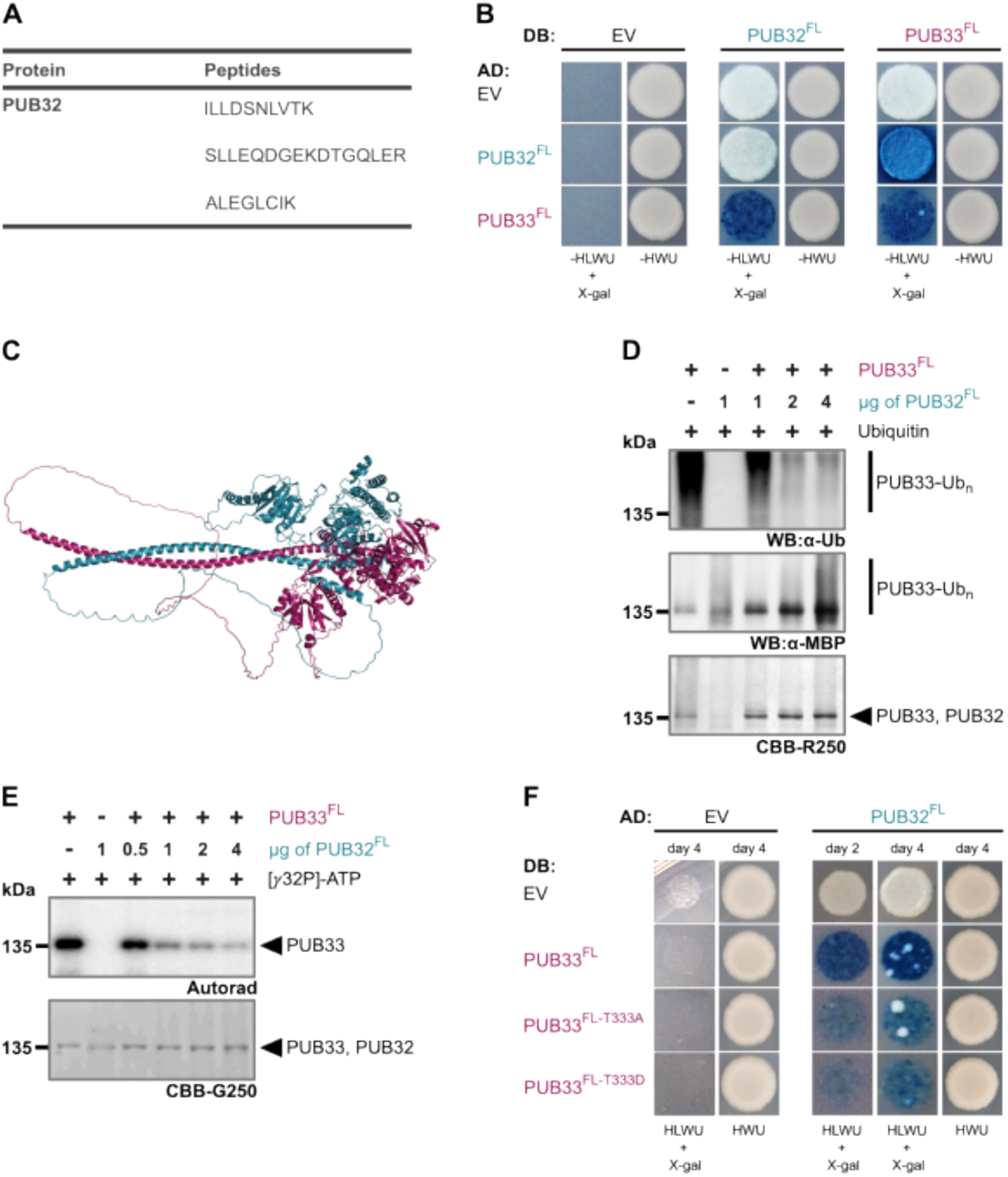
PUB32 inhibits the catalytic activities of PUB33. **(A)** Unique PUB32 peptides identified by mass spectrometry following immunoprecipitation of PUB33-GFP from Col-0/35S:PUB33-GFP line #9-4. Peptides were present in 3/3 PUB33-GFP samples and 0/3 Col-0 samples; see **Supplementary Table S2** for more details. Immunoprecipitation performed by TD; sample processing and analysis performed by MCRG. **(B,F)** Yeast 2-hybrid between PUB32^FL^ and PUB33^FL^ **(B)** and PUB33^FL^ variants **(F)**. EV is the empty vector control. Growth was assessed on SD media lacking His, Leu, Trp, and Uracil (-HLWU) with the addition of X-galactosidase (X-gal) as markers for protein:protein interaction or on SD-HWU as a transformation control. DB indicates constructs translationally fused to the Locus for X-Ray Sensitivity A (LexA) DNA binding domain; AD indicates constructs translationally fused to the B42 transcriptional activation domain. Assays conducted by IK at least three times with similar results. **(C)** Possible configuration of a PUB32:PUB33 homomer, as predicted by AlphaFold2. The colours match the labels in A. Average predicted Local Distance Difference Test (pLDDT) score for the PUB32 monomer is 75.31 and the pLDDT score for the PUB33 monomer is 77.9. The predicted template modeling (pTM) and interface pTM (ipTM) scores for the PUB32:PUB33 heteromer are 0.50 and 0.49, indicating low-to-moderate confidence in both the overall model and the predicted interface. Analysis by TD. **(D)** *In vitro* auto-ubiquitination assays comparing 2 μg His_6_-MBP-PUB33^FL^ in the presence of increasing amounts of MBP-PUB32^FL^. Reaction mixtures include UBA1, UBC8, and ubiquitin. Western blots were probed with anti-Ub and anti-MBP, and protein loading is indicated by post-staining with Coomassie Brilliant Blue (CBB) R250. These assays were repeated more than three times with similar results by TD. **(E)** *In vitro* auto-phosphorylation assays comparing 2 μg His_6_-MBP-PUB33^FL^ in the presence of increasing amounts of MBP-PUB32^FL^. Autoradiography (Autorad) indicates γ-³²P incorporation, and protein loading is indicated by post-staining with CBB G250. Assays conducted at least three times by TD with similar results.

### Genetic manipulation of PUB32 and PUB33 does not impact immune-triggered reactive oxygen species production or defense against *Pseudomonas syringae*

Previous reports suggest that PUB33 may function in immune signaling (He et al. 2019; McLellan et al. 2022; Dong et al. 2023; Choi et al. 2026). In Arabidopsis, sibling plants derived from Salk_139093 were named *pub33-1* and *pub33-2* (referred to here only as *pub33-1*) and reported to display lower flg22-induced ROS and expression of *NHL10* and *FRK1* (Dong et al. 2023). We attempted to isolate homozygous *pub33-1* mutants but were unable to detect the T-DNA insertion in any of the 22 plants we received from the Arabidopsis Biological Resource Center (ABRC). However, we were able to isolate other homozygous alleles, namely Sail_753_D12 (*pub33-3*) and Salk_125263 (*pub33-4*) (**Supplemental Figure S2A**). We also isolated homozygous lines interrupting *PUB32* and *PUB50*, namely Salk_035317 (*pub32-1*), Salk_103257C (*pub32-2*), Salk_113852 (*pub50-1*), and Salk_052482C (*pub50-2*) (**Supplemental Figure S2A**). Quantitative real-time polymerase chain reaction (qPCR) verified loss of expression in these lines (**Supplemental Figure S2B-C**). To assess genetic interactions between *PUB32* and *PUB33*, we also generated a double *pub32-1 pub33-3* mutant. All single and double mutants grew normally in short-day conditions and did not display any obvious developmental defects (**Supplemental Figure S2D**). To our surprise, we did not observe consistent or reproducible aberrations in immune-triggered ROS or susceptibility to the virulent pathogen *Pseudomonas syringae* pathovar *tomato* DC3000 in any of the mutants over a 4-year period (**Supplemental Figure S2E-F**). While we recorded significantly less bacterial growth in *pub33-4* compared to Col-0, we did not observe the same in *pub33-3* (**Supplemental Figure S2F**). These results suggest that PUB33 and PUB32 may not function in innate immune signaling in Arabidopsis, which is contradictory to previously published work reporting lower flg22-induced ROS in the *pub33-1* mutant (Dong et al. 2023). However, as loss of *PUB33* expression was not verified in *pub33-1* (Dong et al. 2023), and only a single allele was tested, it is possible that the mutant does not faithfully report PUB33 function. In addition, the role of PUB33 and its orthologs in flg22-triggered responses is not straightforward. In *N. benthamaiana*, VIGS-mediated knockdown of *NbRLCK-IXb-13* results in lower flg22-induced ROS and gene expression (Dong et al. 2023), and although knockdown of *NbUBK/NbRLCK-IXb-1* results in lower flg22-induced gene expression (He et al. 2019), knockdown of both *NbUBK1/NbRLCK-IXb-1* and *NbRLCK-IXb-2* results in higher flg22-induced ROS (Choi et al. 2026). We therefore find it difficult to make a conclusion about the role of PUB33 and orthologs in innate immune signaling.

### PUB33 ubiquitinates *Ralstonia pseudosolanacearum* effector protein RipV1

Although the biological function of PUB33 and related proteins is not well understood, orthologs of PUB33 interact with effector proteins from highly diverse pathogens. The cyclophilin effector Al106 from *A. lucorum* is able to interact with NbRLCK-1Xb-13 in *N. benthamiana* and AtPUB33 in Arabidopsis (Dong et al. 2023). Conversely, the RXLR effector PiSFI13 from *P. infestans* is only able to interact with StUBK and NbUBK in potato and *N. benthamiana*, but not with AtPUB33 in Arabidopsis (He et al. 2019; McLellan et al. 2022). As Arabidopsis can host *A. lucorum* but not *P. infestans*, it was suggested that PUB33 may be a host-specific target that can be exploited to confer enhanced resistance (McLellan et al. 2022). We recently showed that the NEL effector RipV1 from *R. pseudosolanacearum* can interact with tomato SlRLCK-IXb-1/SlPUB33 and SlRLCK-IXb-2/SlPUB32 via its N-terminal domain (Choi et al. 2026). Since Arabidopsis can host *R. pseudosolanacearum* (Deslandes et al. 1998), it is possible that RipV1 is also capable of interacting with PUB32 and PUB33. To test this, we performed Y2H between DB-RipV1^NT^ (representing the N-terminus of RipV1) and both PUB32^FL^-AD or PUB33^FL^-AD. We found that both can interact with RipV1 (**Figure 6A**). To map the region of interaction, we focused on the PUB33-RipV1 interaction and found that RipV1 interacts with the α-helical domain of PUB33 (**Figure 6A**). We did not observe any difference when we incorporated T333A or T333D mutations into the PUB33^αhelix^-AD construct (**Figure 6A**). Notably, the α-helical domain mediates the interaction between RipV1 and SlRLCK-IXb-1 (Choi et al. 2026), as well as the interaction between ABI5 and PUB35 (Du et al. 2024), suggesting that it serves as a platform for protein interactions.

**Figure 6.**
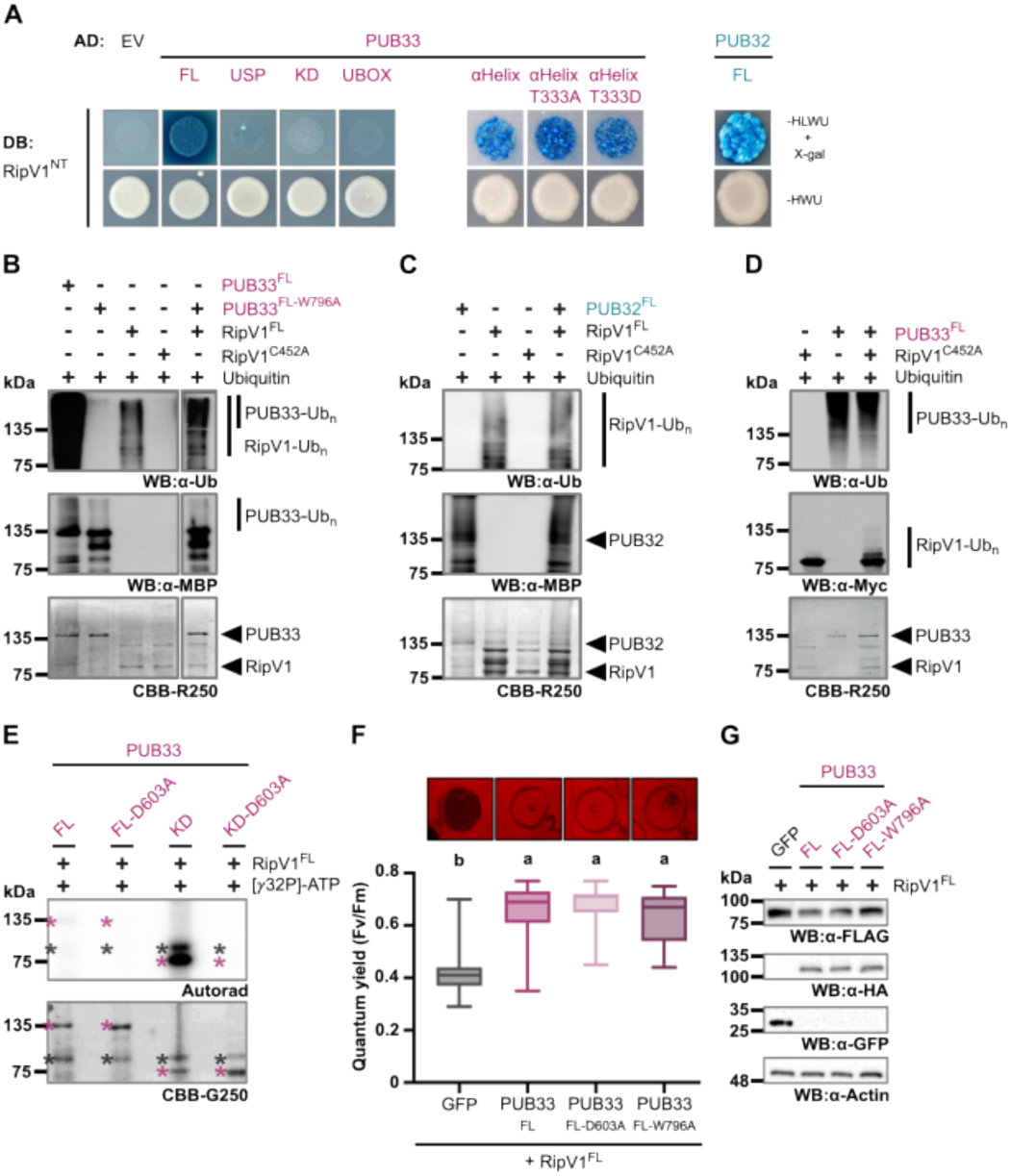
PUB33 ubiquitinates *Ralstonia pseudosolanacearum* effector protein RipV1 and suppresses RipV1-mediated cell death. **(A)** Yeast 2-hybrid between the N-terminal domain of RipV1 (RipV1^NT^) and PUB32^FL^, PUB33^FL^, and the indicated PUB33 variants. EV is the empty vector control. Growth was assessed on SD media lacking His, Leu, Trp, and Uracil (-HLWU) with the addition of X-galactosidase (X-gal) as markers for protein:protein interaction or on SD-HWU as a transformation control. DB indicates constructs translationally fused to the Locus for X-Ray Sensitivity A (LexA) DNA binding domain; AD indicates constructs translationally fused to the B42 transcriptional activation domain. Assays conducted by JC and IK at least three times with similar results. **(B-D)** *In vitro* trans-ubiquitination of PUB33^FL-W796A^ **(B)** and PUB32^FL^ by RipV1^FL^ **(C)**; or RipV1^C452A^ by PUB33^FL^ **(D)**. Reaction mixtures include UBA1, UBC8, and ubiquitin. Western blots were probed with anti-Ub and anti-MBP, and protein loading is indicated by post-staining Coomassie Brilliant Blue (CBB) R250. These assays were repeated more than three times with similar results by TD. **(E)** *In vitro* trans-phosphorylation assay between PUB33^FL^ or PUB33^KD^ and RipV1, compared to the kinase-dead variants containing D603A. Autoradiography (autorad) indicates γ-³²P incorporation, and protein loading is indicated by post-staining with CBB G250. Asterisks indicate proteins of interest. Assays were repeated 3 times by TD. **(F)** Cell death induced by transient expression of RipV1^FL^-FLAG in the presence or absence of PUB33-HA or free GFP in *N. benthamiana*. False color photographs were taken at 3 days post infiltration (dpi); the circles indicate the infiltrated area. Dark areas indicate cell death. Quantum yield of chlorophyll (QY) measurements were also taken at 3 dpi and are summarized in the lower histogram. Values are quantum yield (Fv/Fm) from 3 independent experiments (n=9 leaf discs per experiment). The midline represents the median; interquartile ranges are represented by the boxes, and maximum and minimum values are represented by the whiskers. Significantly different groups are labelled with lower-case letters, based on a one-way analysis of variance (ANOVA) followed by Tukey’s post-hoc test (p<0.0001). These assays were performed by JC and TD more than three times each in independent labs, with similar results. **(G)** Samples from the experiments shown in G were taken at 35 hours post infiltration (hpi) and subjected to western blot. Proteins are detected by anti-FLAG and anti-GFP antibodies; protein loading is indicated by Ponceau S staining. These assays were performed by JC three times with similar results.

As both RipV1 and PUB33 possess E3 ligase activity, we wanted to determine if they are able to ubiquitinate one another. Interestingly, while we previously showed that RipV1 can ubiquitinate SlRLCK-IXb-1 (Choi et al. 2026), we could not clearly detect RipV1-mediated ubiquitination of PUB33^FL-W796A^ or PUB32^FL^ *in vitro* (**Figure 6B,C**). In addition, while we previously could not clearly detect ubiquitination of catalytically inactive RipV1^C452A^ by SlRLCK-IXb-1 (Choi et al. 2026), here we find that PUB33 can ubiquitinate RipV1^C452A^ *in vitro* (**Figure 6D**). Because PUB33 is an active kinase, we also assessed if it can phosphorylate RipV1^FL^ and found that while PUB33^FL^ was not able to phosphorylate RipV1^FL^ *in vitro*, PUB33^KD^ was (**Figure 6E**). Although this suggests that PUB33 may be capable of phosphorylating RipV1, it is important to note that the PUB33^KD^ variant does not contain the α-helical domain which is the site of their interaction, so these data are difficult to interpret. We propose that it may be possible for PUB33 to phosphorylate RipV1, however we currently do not have enough data to robustly support this hypothesis.

Like many pathogen effectors, RipV1 induces cell death when expressed in *N. benthamiana* (Cheng et al. 2024; Choi et al. 2026). However, as RipV1-induced cell death cannot be suppressed by blocking canonical NLR signaling, it is likely a symptom of disease rather than immune activation (Choi et al. 2026). Over-expression of *SlRLCK-IXb-1/SlPUB33* suppresses RipV1-induced cell death, suggesting that SlRLCK-IXb-1/SlPUB33 directly inhibits RipV1 function. To test if PUB33^FL^ is capable of suppressing RipV1-induced cell death, we co-expressed both proteins in *N. benthamiana* and found that this was indeed the case (**Figure 6F**). Furthermore, western blot analysis indicates that the level of RipV1 is drastically reduced in the presence of PUB33^FL^ at 35 hours post infiltration (hpi) (**Figure 6G**). Because PUB33 can ubiquitinate RipV1 *in vitro* (**Figure 6D**), we hypothesized that this may be due to PUB33-mediated ubiquitination and subsequent degradation of RipV1 via the 26S proteasome. Protein accumulation of RipV1 was less impacted when co-expressed with either AtPUB33^FL-D603A^ or PUB33^FL-W796A^ (**Figure 6G**), corroborating that hypothesis. However, we also found that both PUB33^FL-D603A^ and AtPUB33^FL-W796A^ are able to suppress RipV1-mediated cell death similarly to PUB33^FL^ (**Figure 6F**), indicating that a reduction in RipV1 accumulation cannot solely explain the suppression of cell death. This uncoupling of RipV1 degradation from suppression of cell death suggests that PUB33 interferes with RipV1 function through additional, ubiquitination-independent mechanisms. The ability of SlRLCK-IXb-1 to suppress RipV1-induced cell death without detectable ubiquitination of RipV1 (Choi et al. 2026) further supports a model in which suppression occurs at the level of effector activity rather than effector abundance. Together, these observations imply that AtPUB33 and SlRLCK-IXb-1 act as a negative regulator of RipV1 virulence, potentially by sequestering RipV1, altering its subcellular localization, or preventing its interaction with host targets. Importantly, the lack of immune signaling defects in *pub32*, *pub33*, and *pub32 pub33* mutants argues against a general role for PUB33 in canonical immune pathways and instead points to a RipV1-specific function. Given that RipV1 also interacts with PUB32, a highly similar but non-catalytic modulator of PUB33 function, it is plausible that PUB32 represents the *bona fide* virulence target of RipV1, with PUB33 acting as a functional decoy. In this model, PUB33 could trap the effector, attenuating its activity through both ubiquitination dependent and independent mechanisms. Further work is needed to test this hypothesis.

## Methods

### Data mining and phylogenetics

Protein sequences of Arabidopsis PUB-IV/RLCK-IXb family members were retrieved from The Arabidopsis Information Resource (TAIR) (Berardini et al. 2015) and the Plant Transcription Factor and Protein Kinase Identifier and Classifier database (iTAK) (Zheng et al. 2016). Multiple sequence alignments were generated using the MUSCLE algorithm implemented in Molecular Evolutionary Genetics Analysis (MEGA X) (Kumar et al. 2018). Phylogenetic trees were constructed in MEGA X using the maximum likelihood method with 100 bootstrap replicates to assess branch support. Resulting trees were annotated and visualized with the Interactive Tree of Life (iTOL) platform (Letunic and Bork 2024). To examine expression profiles, transcriptome data of RLCK-IXb genes across different developmental stages were obtained from the Arabidopsis Electronic Fluorescent Pictograph (eFP) Browser (Winter et al. 2007), and the processed data were used to generate heat maps in GraphPad Prism (version 10.0.0 for macOS, GraphPad Software, Boston, Massachusetts USA, https://www.graphpad.com). For structural and functional characterization, conserved motifs and domains were identified using the Pfam database (EMBL-EBI Pfam 34.0) (Mistry et al. 2021), and predicted three-dimensional protein structures were retrieved from the AlphaFold Protein Structure Database (Jumper et al. 2021) and further processed in PyMOL (The PyMOL Molecular Graphics System, Version 2.1 Schrödinger, LLC) or ChimeraX (Pettersen et al. 2021). Gene organization and domain architecture were illustrated using Illustrator for Biological Sciences (IBS) version 2.0 (Xie et al. 2022).

### Germplasm and plant growth conditions

*Arabidopsis thaliana* and *Nicotiana benthamiana* plants were grown in the Queen’s Phytotron Facility. For aseptic growth, *A. thaliana* seeds were surface-sterilized using a 40% bleach solution for 15 minutes and sown onto petri plates with 0.5x Murashige and Skoog (MS) media containing 0.8% agar. Afterward, they were subjected to cold stratification in darkness at 4°C for 3-5 days before being exposed to light. For plants grown in soil, seeds were directly sown onto potting soil (Sungro Sunshine Mix 1 or PRO-MIX FPX AGTIV STIMULATE fine plug growing medium). Seedlings were later transplanted either individually into 3” pots or six plants per 8” pot two weeks after sowing. Controlled growth chambers (BioChambers and Conviron) were utilized for plant cultivation, maintaining a 10-hour light and 14-hour dark cycle at 22°C, with 30% relative humidity and a light intensity of 150 µE m^2^ s^-1^. Plants were watered from the top as needed, typically every other day, and fertilized every two weeks with a solution containing 1.5 g/L of 20:20:20 N:P:K. *N. benthamiana* plant cultivation followed a similar procedure: seeds were sown on potting soil and transplanted individually, but were grown in a dedicated growth chamber (Conviron) with a 16-hour light and 8-hour dark cycle, receiving weekly fertilization. Additionally, mite bags containing *Amblyseius swirskii* (Koppert) or *Neoseiulus cucumeris* (Koppert) were introduced bi-weekly to each plant tray to prevent pest infestations.

Information on all germplasm used in this study is outlined in detail in **Supplemental Table S2**. Segregating T-DNA insertion lines were obtained from the Arabidopsis Biological Resource Centre (ABRC) and genotyped using gene-specific primers in standard polymerase chain reactions (PCR). To determine gene expression, we extracted RNA using the Aurum Total RNA Mini Kit (BioRad), according to the manufacturer’s instructions. Reverse transcription (RT) was achieved using Superscript III (Invitrogen), according to the manufacturer’s instructions. Quantitative RT-PCR was performed using Sso Advanced Universal SYBR Green Supermix (BioRad) according to the manufacturer’s directions, on a CFX96 Touch Real-Time PCR Detection System (BioRad), using primers specific to each gene as outlined in **Supplemental Table S2**. Reference genes used were *ACTIN2* and *PP2A3* (Czechowski et al., 2005). Our analysis included three technical replicates from three biological samples per genotype (n=9); each biological sample included leaf tissue from different plants. Primer efficiencies were >100% and similar for all pairs and melting curve analyses indicated one amplicon per pair. Higher-order mutants were generated by crossing and genotyping to homozygosity in the F_2_ and F_3_ generations. Transgenic plants were generated by floral dip with *Agrobacterium tumefaciens* GV3101 carrying a suitable binary vector for selection, as previously described (Clough and Bent 1998). Independent lines were followed through four generations and genotyped to homozygosity based on 100% resistance to the selection marker in the T_3_ generation.

### Molecular cloning

Detailed information about all vectors used and generated in this study can be found in **Supplemental Table S2.** Gateway-compatible pENTR clones were synthesized by Twist BioSciences (San Francisco, USA), assembled via Gibson assembly (New England Biolabs) using PCR fragments amplified with Q5 DNA polymerase, or obtained from the ABRC (Feke et al. 2019). Recombination of entry vectors into Gateway-compatible binary vectors pK7FWG2 (Karimi et al. 2002) was achieved using Gateway LR Clonase II (Invitrogen) according to the manufacturer’s instructions. For recombinant protein production in *Escherichia coli*, complete pET28a+ constructs were synthesized by Twist BioSciences (San Francisco, USA). In some cases, gene coding regions were either synthesized as linear fragments or PCR-amplified using Q5 DNA polymerase (New England Biolabs) and cloned into pMAL-c2x (Walker et al. 2010) following the manufacturer’s protocols, employing construct-specific endonucleases and T4 DNA ligase (New England BioLabs). Additionally, pDEST-HisMBP (Nallamsetty et al. 2005) was used for Gateway cloning to generate recombinant proteins in *E. coli* from pENTR-D-TOPO clones, as listed in **Supplemental Table S2**. Successful assembly of all plasmids was verified using whole-plasmid sequencing (Plasmidsaurus, Eugene OR, USA). Details of the *E. coli* expression constructs and plant expression clones for RipV1 are provided in **Supplemental Table S2**.

### Immune and growth assays

Immune-triggered production of apoplastic reactive oxygen species (ROS) and infection with *Pseudomonas syringae* pv. *tomato* DC3000 were performed as described previously (Monaghan et al. 2014; Bredow et al. 2019; Gonçalves Dias et al. 2024). Immunogenic flg22, elf18, and AtPep1 peptides were synthesized by EZ Biotech (Indiana USA).

### Cell death assay, quantification and testing protein accumulation

Cell death assays were performed in the Segonzac lab following a protocol described previously (Choi et al. 2026), and in the Monaghan lab using the following modifications. Cultures of *Agrobacterium tumefaciens* GV3101 adjusted to an OD_600_ of 0.25 and carrying appropriate plasmids listed in **Supplemental Table S2** were infiltrated into fully expanded leaves of 3- to 4-week-old *N. benthamiana* plants together with the viral suppressor P19. At four days post-infiltration, leaves (attached or detached from the plant) were imaged using the NightShade LB985 In Vivo Plant Imaging System (Berthold Technologies) in luminescence mode to measure delayed fluorescence (DF). DF, which reflects photosynthetic activity, arises from the re-excitation of chlorophyll molecules in Photosystem II, was quantified following the manufacturer’s protocol. Images were analysed using IndiGO™ software (Berthold Technologies), and fluorescence intensity was expressed as counts per second per unit area (cps/area). To assess protein accumulation, leaf discs were harvested at 35 hpi followed by western blotting.

### Recombinant protein purification and *in vitro* kinase and ubiquitination assays

All proteins were expressed and purified from *E. coli* strain BL21 cells as recently described (Gonçalves Dias et al. 2024), using the constructs outlined in **Supplemental Table S2**. *In vitro* kinase assays using γ32P-ATP were performed as recently described (Gonçalves Dias et al. 2024), using 2 μg kinase in a buffer containing 50 mM Tris-HCl (pH 8.0), 25 mM MgCl_2_, 25 mM MnCl_2_, 5 mM DTT, 5 μM ATP, and 0.5-2 μCi γ32P-ATP. Histone 3S from calf thymus was used as a universal substrate in some reactions (Sigma Aldrich H5505). All reactions were incubated for 30 minutes or 45 minutes at 30°C. Reactions were stopped by adding 5x Laemmli buffer and heating at 80°C for 5 min. Proteins were separated by 8% SDS-PAGE. The gels were sandwiched between two sheets of transparency film, exposed to a storage phosphor screen (Molecular Dynamics) overnight, and visualized using Typhoon 8600 Imager (Molecular Dynamics/Amersham), Typhoon FLA 9500 imager (Cytiva/Amersham), Amersham Typhoon (Cytiva), or Sapphire FL Biomolecular Imager (Azure Biosystems). Gels were post-stained with Coomassie Brilliant Blue (CBB) R-250 (MP Biomedicals) or SimplyBlue SafeStain (Invitrogen; CBB G-250) and scanned.

*In vitro* auto-ubiquitination assays were performed as described by Dou *et al*. (2024). For auto-ubiquitination assays, 2 μg of recombinant E3 ligase and 0.075 μg of HA-tagged ubiquitin (R&D Systems; U-110) were incubated in 40 μL of reaction buffer containing 25 mM Tris-HCl (pH 7.4), 5 mM MgCl₂, 25 mM KCl, 0.33 mM DTT, 1.5 mM ATP, 200 ng E1 (UBA1; AT2G30110), and 450 ng E2 (UBC8; AT5G41700) for the indicated times. For trans-ubiquitination assays, 4 μg of substrate protein was added, and reactions were performed in 60 μL of the same buffer. Reactions were terminated by adding 5x Laemmli Sample Buffer, followed by heating at 80°C for 5 minutes.

### Immunoblotting

Proteins were separated on 8% or 10% SDS–PAGE gels, transferred to polyvinylidene difluoride (PVDF) membranes (BioRad), and blocked with 5% non-fat milk in Tris-buffered saline containing 0.05–0.1% Tween-20. The following antibodies were used, as indicated in the figure captions: anti-Ub (Cell Signaling Technology; P4D1); anti-MBP (New England Biolabs; E8032S); anti-HA (Roche; 12013819001); anti-Myc (Cell Signaling Technology; 9B11); anti-GFP (Roche; 1814460001 or ChromoTek; 3h9-150); anti-FLAG-HRP (Sigma; A8592); anti-mouse (Sigma, A0168); anti-mouse (LiCOR Biosciences; IRDye 800CW); anti-rat (Sigma; A5795). For chemiluminescence, membranes were incubated with ECL Clarity Substrate (BioRad) and imaged with a ChemiDoc Touch Imaging System (BioRad). For fluorescence detection, blots were scanned using a Odessy®-XF imaging system (LiCORbio^TM^). After imaging, blots were stained with Coomassie Brilliant Blue R-250 (MP Biomedicals) as a loading control.

### Yeast 2-hybrid

The yeast strain EGY48 (mating type α) carrying the reporter plasmid pSH18-34 was transformed with the bait constructs cloned into the pLexA vector, while RFY206 (mating type a) was transformed with prey contrsucts cloned into the pB42-AD vector, using the Frozen-EZ Yeast Transformation II Kit (Zymo Research; T2001). All constructs used in this study are listed in **Supplemental Table S2**. Transformants harboring bait constructs were selected on synthetic defined (SD) dropout medium lacking uracil and histidine (SD/-His/-Ura; Clontech; 630422), whereas transformants carrying prey constructs were selected on SD medium lacking tryptophan (SD/-Trp; Clontech; 630413). For yeast mating, bait- and prey-expressing yeast strains were co-cultured in 200 µl of yeast extract peptone dextrose (YPD) broth (Difco; 242820) and incubated with shaking overnight at 30 °C. Diploid cells were selected on SD medium lacking histidine, tryptophan, and uracil (SD/-His/-Trp/-Ura; Clontech; 630424). To assess protein-protein interactions, diploid yeast cells were plated onto SD medium lacking histidine, leucine, tryptophan, and uracil (SD/-His/-Leu/-Trp/-Ura; Clontech; 630425) medium supplemented with raffinose (Sigma-Aldrich; 83400), galactose (Sigma-Aldrich; G0750), and 5-bromo-4-chloro-3-indolyl β-D-galactopyranoside (X-gal; LPS solution; XGAL5000) to induce prey protein expression and monitor reporter gene activation. Interaction-dependent activation of the *LEU2* and *lacZ* reporter genes were evaluated based on yeast growth and blue color development, respectively. Plates were incubated at 30 °C for 2-4 days before being photographed.

### Immunoprecipitation

Ten-day-old *Arabidopsis* seedlings stably expressing GFP-tagged proteins were harvested, ground in liquid nitrogen, and resuspended in plant protein extraction buffer (0.5 M Tris-HCl pH 7.5, 0.15 M NaCl, 10% glycerol, 2 mM NaF, 1.5 mM activated Na₃VO₄, 1 mM DTT, 1% Igepal, 1 mM PMSF) supplemented with 1% protease inhibitor cocktail (Sigma-Aldrich; P9599). Extracts were incubated with GFP-Trap agarose beads (Chromotek; gta) for 2 hours at 4°C. Beads were collected by centrifugation and washed three times with wash buffer (1x TBS, 1 mM DTT, 1% Igepal) supplemented with 1% protease inhibitor cocktail (Sigma-Aldrich; P9599). Bound proteins were eluted with 5x Laemmli sample buffer and denatured at 80°C for 5 min before use in SDS-PAGE and Western blot analysis.

### Mass spectrometry and phosphoproteomics

To map PUB33-mediated phosphorylation sites, *in vitro* kinase assays were performed using 2 μg of His_6_-MBP-PUB33-FL or His_6_-MBP-PUB33-FL-D603A in a buffer containing 25 mM Tris–HCl (pH 8.0), 10 mM MgCl_2_, 100 μM MnCl_2_, 1 mM DTT, 1 mM phenylmethylsulfonyl fluoride (PMSF), and 100 μM ATP. All reactions were incubated for 60 min at 30 °C and stopped by adding 5x Laemmli sample buffer and heating at 80 °C for 5 min. Protein separation was carried out using 10% Mini-PROTEAN TGX Precast Gels (BioRad; 456-1036), followed by post-staining with SimplyBlue SafeStain (Invitrogen; CBB G-250). Bands corresponding to the size of His_6_-MBP-PUB33-FL or His_6_-MBP-PUB33-FL-D603A were excised using sterile razors and washed with ethanol and sterile ddH_2_O. Subsequent steps and phosphoproteomics were performed exactly as described in (Gonçalves Dias et al. 2024). To identify PUB33-associated proteins, *Arabidopsis* Col-0 and Col-0/35S:PUB33-GFP homozygous line 9-4 seedlings were processed for immunoprecipitation as described above. Denatured protein extracts were separated on 10% Mini-PROTEAN TGX Precast Gels (BioRad; 456-1036). When proteins had just entered the resolving gel from the stacking gel, electrophoresis was stopped, and the gels were post-stained with SimplyBlue SafeStain (Invitrogen, CBB G-250). Bands corresponding to total protein were excised using sterile scalpels and washed with ethanol and sterile ddH_2_O. Phosphoproteomic analyses were performed as previously described (Gonçalves Dias et al. 2024) without deviation. PUB33-GFP data was acquired using direct data independent acquisition (dDIA) as previously described (Mehta et al. 2022), without deviation, using a 68 minute sigmoidal gradient. Acquired dDIA mass spectrometry data was analyzed using Spectronaut v19 (Biognosys AG) under default settings. Data were searched using the Arabidopsis thaliana ARAPORT 11 proteome accessed through Phytozome (https://phytozome-next.jgi.doe.gov/info/Athaliana_Araport11). Search parameters included the following: trypsin digest allowing two missed cleavages, fixed modifications (carbamidomethyl [C]), variable modifications (oxidation [M]), a peptide spectrum match, and peptide and protein false discovery threshold of 0.01. Significantly changing differentially abundant proteins were determined and corrected for multiple comparisons (Bonferroni-corrected *p* value < 0.05; *q* value). Protein interactome data was subsequently assessed using the acquired MS1 quantification data in combination with the Significance Analysis of INTeractome software package (SAINTq) (Teo et al. 2016); an interaction significance threshold of q < 0.05 was used.

### Software

Multiple sequence alignments and phylogenetic analyses were performed using Molecular Evolutionary Genetics Analysis (MEGA X) (Kumar et al. 2018). Data processing and statistical analyses were carried out in GraphPad Prism (version 10.0.0 for macOS, GraphPad Software, Boston, Massachusetts USA, https://www.graphpad.com). AlphaFold-predicted protein structures were visualized and refined using PyMOL (The PyMOL Molecular Graphics System, Version 2.1 Schrödinger, LLC) or ChimeraX (Pettersen et al. 2021). Western blot and post-stained gel images were processed using ImageJ (Schneider et al. 2012). Final figures were assembled in Inkscape v1.4.2 (Inkscape Project. (2020). Inkscape. Retrieved from https://inkscape.org).

## Data Availability

All raw proteomic data have been uploaded to ProteomeXchange (http://www.proteomexchange.org/) via the Proteomics IDEntification Database (PRIDE; https://www.ebi.ac.uk/pride/). Project Accession: PXD073806. Username. Password: pjIqhpoygoEX

## Author Contributions

TD performed the majority of the work and co-wrote the paper with JM. JC, IK, VNM, and NSK generated materials, performed experiments, and analyzed results. JC and IK performed RipV1-induced cell death assays and all Y2H, supervised by CS. MCRG performed phosphoproteomics and PUB33-IP proteomics; NH performed statistics on the PUB33-IP dataset; MCRG and NH were supervised by RGU. TD worked in the lab of MT for 3 months to learn various techniques in PUB biochemistry. Individual credits are included wherever possible in the figure captions and table legends. JM designed the project, guided the work, supervised TD and VNM, secured funding, and wrote the paper with input from all authors.

## Acknowledgements

We thank all members of the Monaghan Lab for their comments on this manuscript and for their commitment to fostering a welcoming and collaborative research environment. We thank Emily DeSousa and Nicholas Smith for contributing to pilot work on this project as undergraduates; Saeid Mobini for managing the Queen’s University Phytotron Facility; Xiaojing Yang and Neil Renwick for their assistance with phosphorimaging at Queen’s University; and Jack Moore for technical assistance and maintenance of the University of Alberta Mass Spectrometry and Proteomics Facility. We thank Carla Brillada and Bushra Saeed from the Trujillo Lab for their support in establishing the ubiquitination assays. Queen’s University is situated on the territory of the Haudenosaunee and Anishinaabek and we are grateful to live, work, and play on these lands.

## Supplemental Data

Supplemental Table S1. PUB33-associated proteins identified by mass spectrometry.

*Provided as a separate file*.

Supplemental Table S2. Germplasm, clones, and oligonucleotides used in this study.

*Provided as a separate file*.

**Supplemental Figure S1.**
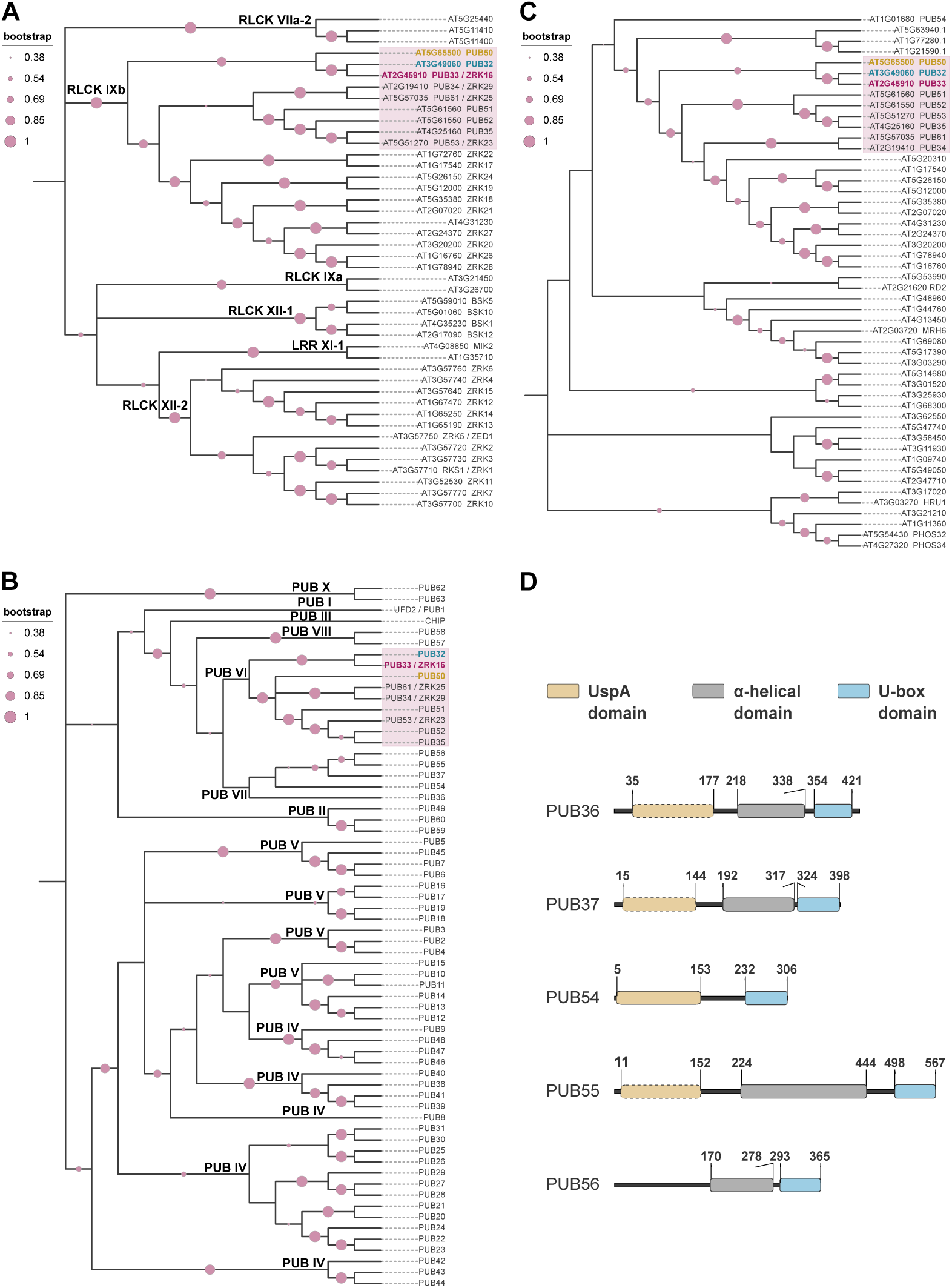
Phylogenetic analysis of RLCK-IXb proteins. Maximum likelihood phylogenetic trees showing the relatedness of RLCK-IXb proteins to their closest homologs within the RLCK **(A)**, PUB **(B)**, and UspA **(C)** families, supported by 100 bootstraps. Well-defined subgroups are labeled for the RLCK proteins as per (Shiu and Bleecker 2003) and PUB proteins as per (Trenner et al. 2022). **(D)** Protein domain architecture for PUB-VII proteins, which contain a UspA domain, α-helical domain, and U-box domain, but lack the RLCK domain found in PUB-VI/RLCK-IXb proteins. Amino acid boundaries are labelled and based on Pfam analysis and manual inspection of AlphaFold predictions. Analysis by TD.

**Supplemental Figure S2.**
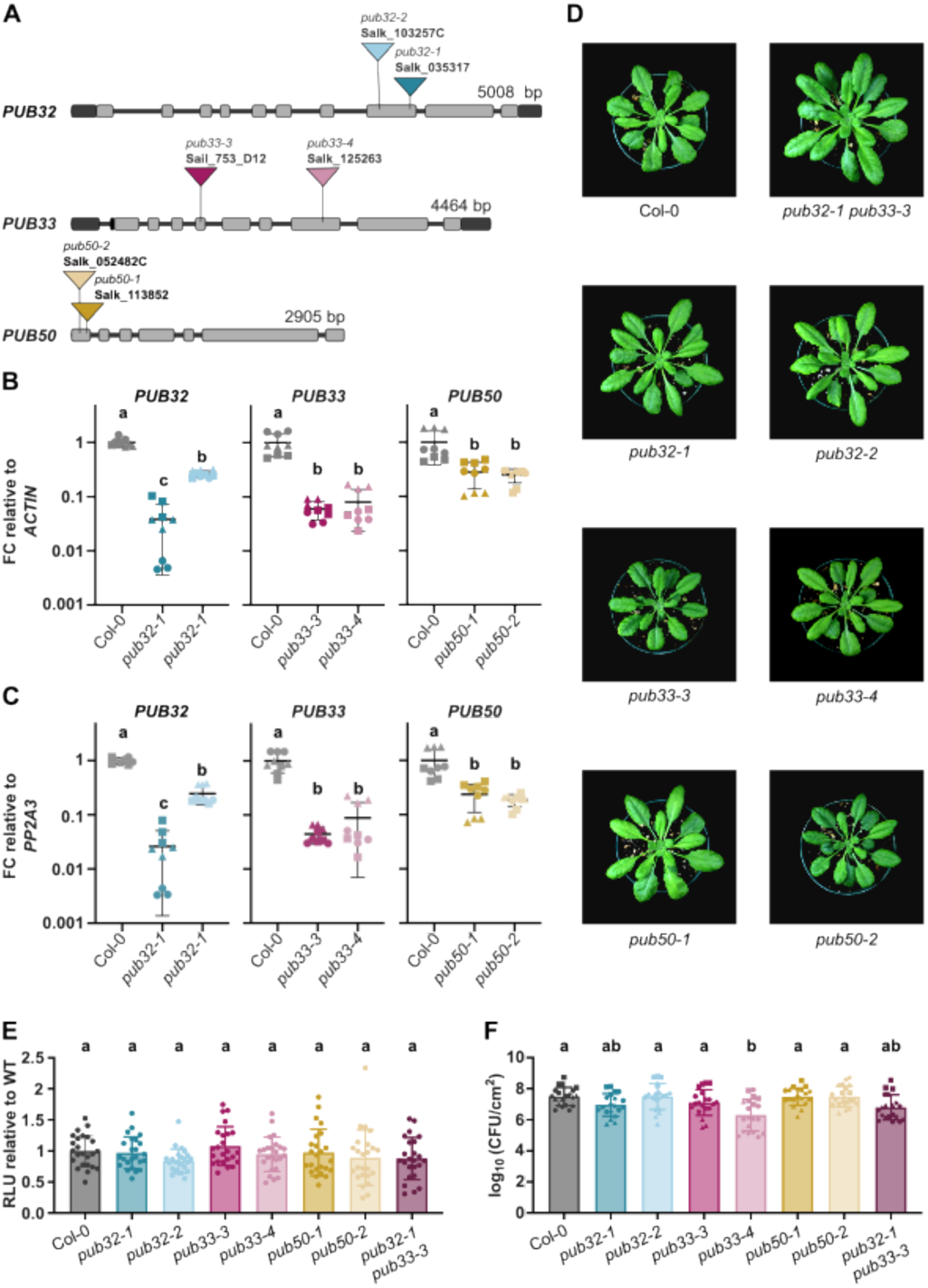
Analysis of Arabidopsis loss-of-function lines in *PUB32, PUB33*, and *PUB50*. **(A)** Schematic representation, drawn to scale, of Arabidopsis RLCK-IXb genes, indicating introns (lines), translated regions (light gray boxes), untranslated regions (dark grey boxes), and the location of T-DNA insertion alleles. Genomic information was retrieved from The Arabidopsis Information Resource (TAIR) and plants were genotyped to homozygosity by TD as outlined in **Supplemental Table S2**. **(B-C)** Quantitative real-time qRT-PCR of *PUB32*, *PUB33*, or *PUB50*, relative to *ACTIN* **(B)** and *PP2A3* **(C)** in the indicated genotypes, presented as fold-change (FC) relative to Col-0. FC values are plotted on a log₁₀-scaled Y-axis to improve visualization of differences among genotypes. FC values themselves were not transformed. Values represent 3 technical replicates for 3 biological samples (differentiated by circle, triangle, or squares); the line represents the mean; and the whiskers represent standard error of the mean. Each biological sample represents multiple leaves from different plants of the same genotype. Lower-case letters indicate statistically significant groups determined by a one-way ANOVA followed by Tukey’s post-hoc test (p<0.05). All data collected by VNM. Primers for genotyping and qRT-PCR are provided in **Supplemental Table S2**. **(D)** Photographs of representative plants of each genotype after 5 weeks of growth on soil under short-day conditions. Photographs taken by TD. **(E)** Reactive oxygen species production measured in relative light units (RLUs) after treatment with 200 nM flg22. Values represent RLU relative to the mean value of Col-0 (n=6) for each respective experiment; whiskers indicate standard deviation. Results from four independent assays are plotted together. None of the genotypes are significantly different from each other, as determined by a one-way ANOVA followed by Tukey’s post-hoc test. **(F)** Growth of *Pseudomonas syringae* pv. *tomato* (*P.s.t.*) isolate DC3000 3 days after syringe-inoculation. Data from five independent biological replicates are plotted together, denoted by squares, circles, triangles, diamonds, and pentagons. Values are colony forming units (cfu) per leaf area (cm^2^) from 4-5 samples per genotype (each sample contains 3 leaf discs from 3 different infected plants); whiskers indicate standard deviation. Lower case letters indicate statistically significant groups, as determined by a one-way ANOVA followed by Tukey’s post-hoc test (p<0.05). Data collected by TD.

